# DRAFT: graph-recurrent learning of dynamic gene regulatory networks along single-cell multi-omics trajectories

**DOI:** 10.64898/2026.11.17.767145

**Authors:** Zehua Zhang, Pallavi Santhi Sekhar, Patrick van Nierop y Sanchez, Ingrid Lohmann, Britta Velten

## Abstract

Single-cell multi-omics data offer an opportunity to resolve how gene regulatory networks (GRNs) are rewired during cell-fate transitions, but most existing methods infer static snapshots, ignore temporal dependencies or assume linear regulation. Here we present DRAFT, a deep learning framework that combines graph attention with recurrent temporal memory to infer dynamic GRNs from time-resolved single-cell multi-omics data. On simulated data, DRAFT outperforms existing methods in accuracy and temporal smoothness. In mouse cellular reprogramming and *Drosophila* adult spermatogenesis, it recovers interactions supported by independent ChIP-seq, perturbation and Hi-C data, together with known stage-specific regulators. By inferring regulatory activity from dynamic network structure rather than expression alone, DRAFT identifies regulators whose activity precedes their transcriptional peak, including the candidate early germline regulator *Mondo*. Germline-specific *Mondo* knockdown alters germline stem cell number and expression of the predicted targets *kuz* and *mamo*. DRAFT also enables nonlinear alignment of co-developing lineages and reveals stage-specific cell–cell communication.

## Introduction

Gene regulatory networks (GRNs) reflect the interaction mechanisms among transcription factors (TFs), regulatory elements (REs), and target genes (TGs), and they provide a valuable basis to understand the regulatory rules underlying development, cell identity and cell state transitions^1^. In biological processes such as development, GRNs typically change over time due to changes in the chromatin landscape or context-dependent presence of co-factors^2,3^, leading to cell-type-specific gene expression profiles. Single-cell multi-omics sequencing, more precisely, single-cell RNA sequencing (scRNA-seq) and single-cell Assay for Transposase-Accessible Chromatin using sequencing (scATAC-seq), measures gene expression profiles and chromatin accessibility states at single-cell resolution along biological processes and organizes them as single-cell trajectories^2^. This provides unprecedented resolution for analyzing cell-state transitions. However, to translate these data, which record transient transcriptional changes, into an understanding of the dynamics of gene expression mechanisms, computational methods are still needed to construct dynamic GRNs over time.

Although numerous methods infer GRNs from multi-omics data^1,4–9^, computational methods for reconstructing dynamic GRNs along single-cell multi-omics trajectories remain limited^10–14^, mainly because of the following difficulties: (1) Temporal dependencies: Single-cell trajectories capture the temporal relationships between cell states, suggesting that GRN inference should leverage not only measurements from individual states but also information from neighboring states along the trajectory. (2) Temporal smoothness: Regulatory networks along a continuous biological process are expected to exhibit gradual transitions, requiring adequate model regularization. (3) Nonlinear regulation: TG expression is governed by complex, potentially nonlinear mechanisms^5^ involving TF expression and RE accessibility, which are challenging to capture with simplified linear models. Existing approaches (e.g., SCENIC+^1^, scMTNI^13^, Dictys^11^, CellOracle^10^, scTIE^14^, scDoRI^12^) focus on static network reconstruction, adopt simplified linear regulatory assumptions, or incorporate temporal information only through smoothing or regularization (**Supplementary Table 1**). However, jointly modeling nonlinear regulatory interactions, temporal dependencies, and temporal smoothness within a unified multi-omics framework remains challenging.

Deep learning offers tools to address these challenges. Graph neural networks (GNNs), such as graph convolutional networks (GCNs)^15^, graph attention networks (GATs^16^) and GraphSAGE^17^, nonlinearly aggregate information over each node’s neighborhood and have been used to infer static regulatory networks from priors and multi-omics data (e.g., GLUE^6^, DeepMAPS^18^). Meanwhile, recurrent networks (e.g., LSTM^19^, GRU^20^) transmit and filter historical information across adjacent stages and have also been used to model temporal dependencies in dynamic GRN inference from RNA-seq time series (Marlene^21^).

Based on these advances, we propose **DRAFT** (**D**ynamic Gene Regulatory Network Inference with Graph-**R**ecurrent **A**ttention **f**rom Single-cell Multi-omics **T**rajectories), a deep learning framework for constructing dynamic GRNs from single-cell multi-omics trajectories. DRAFT supports paired RNA–ATAC profiles from the same cells and unpaired profiles measured in separate cells. By combining graph attention with recurrent temporal memory, DRAFT integrates regulatory priors and molecular measurements while explicitly modeling dependencies between stages, thereby producing dynamic GRNs that gradually change over time. For paired data, DRAFT infers enhancer-driven GRNs (eGRNs), in which TFs regulate TGs through REs; we summarize key GRN subnetworks as enhancer-driven regulons (eRegulons), each comprising a TF, its putatively bound REs and the TGs linked to those REs, and examine the main regulatory relationships with cluster and topic analyses.

To evaluate DRAFT, we benchmark accuracy and temporal smoothness in simulations with ground-truth GRNs. We next apply DRAFT to unpaired mouse cellular reprogramming and paired *Drosophila* testis data. We assess the inferred networks against independent ChIP-seq, perturbation and Hi-C data. Further, DRAFT predicts regulators whose activity precedes their transcriptional peak. We validate a predicted early germline role for *Mondo* through germline-specific knockdown and measure transcript levels of its candidate targets, *kuz* and *mamo*. Finally, we extend the analysis to nonlinear alignment and ligand–receptor timing across co-developing lineages.

## Results

### DRAFT combines regulatory graph learning and temporal memory for dynamic GRN inference

DRAFT provides a graph learning framework to infer stage-specific GRNs from single-cell multi-omics trajectories (**Fig. 1a**). The model takes as input scRNA and scATAC data measured on the same (“paired”) or separate (“unpaired”) cells along a linear or branching trajectory of discrete stages, such as sampling times, cell clusters or pseudotime intervals. As a starting point the model first constructs a static prior graph representing putative regulatory interactions. For paired data, this graph includes TFs, TGs and REs with edges based on known motif binding sites (TF-RE links)^1,4^ and genomic proximity (RE–TG links). For unpaired data, accessible motif-containing regions are used to construct a prior graph containing only TF and TG nodes. For each stage, the prior graph is then equipped with stage-specific RNA measurements and, in the case of paired data, chromatin accessibility data as node features (**Fig. 1a, Methods**).

**Fig. 1.**
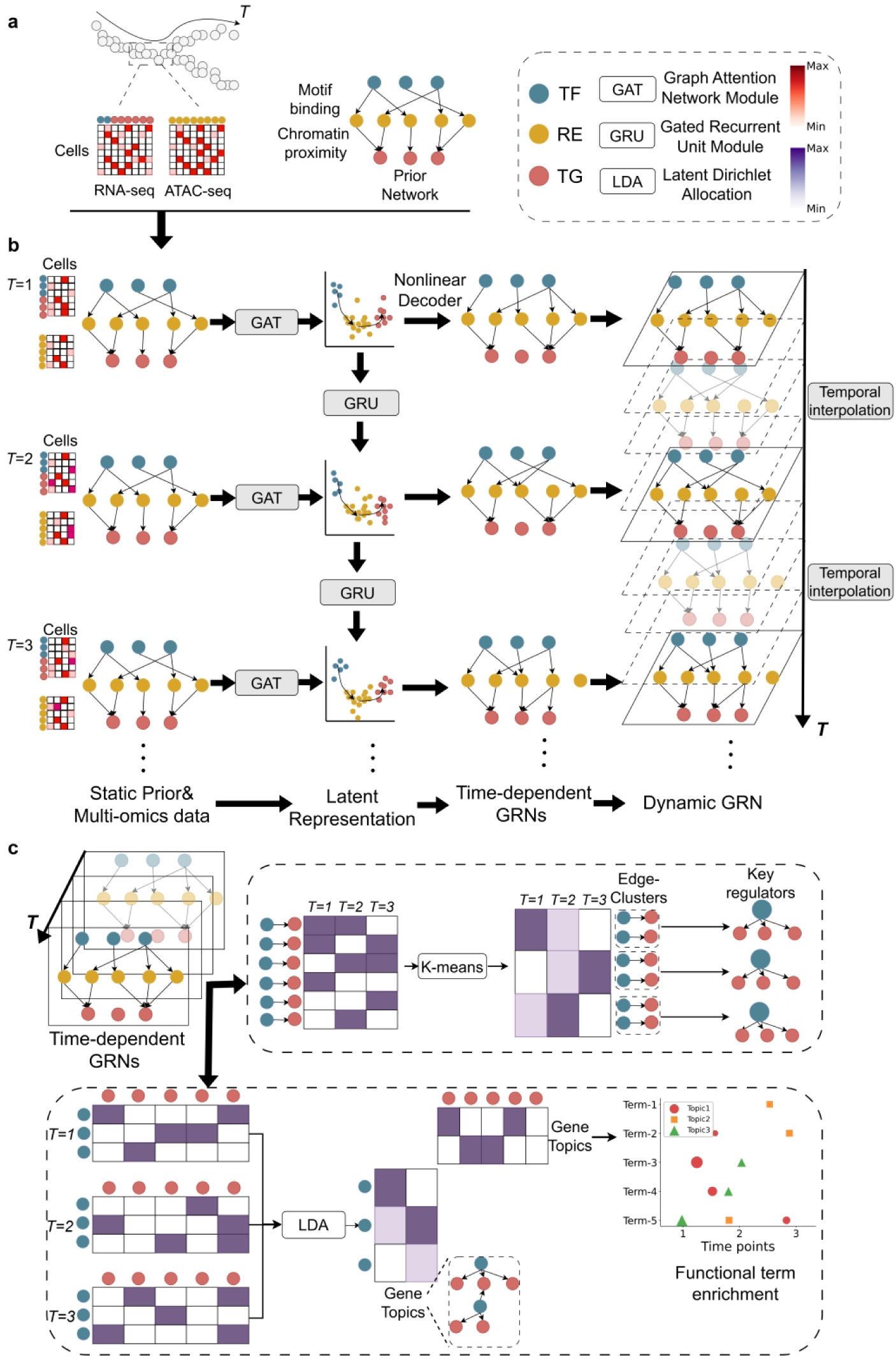
DRAFT model overview. **a**, The DRAFT model first constructs a static prior graph *G_0_* calculated by motif binding (TF-RE relationships) and genomic distance (RE-TG relationships). Stage-specific gene expression and chromatin accessibility data are input into the model as node feature vectors. **b,** Core modules of DRAFT include a Graph Attention Network (GAT) which aggregates information at each stage and a Gated Recurrent Unit (GRU) which links stages to model temporal dependency. These modules are jointly trained within a unified framework to obtain stage-specific latent representations for transcription factors (TFs), regulatory elements (REs), and target genes (TGs). Similarity in the latent space is used to score regulatory strengths, based on which the model reconstructs and outputs stage-specific regulatory networks and eRegulons. Then, DRAFT uses a Gaussian kernel interpolation to interpolate unseen stages and obtain continuously varying GRNs. **c,** To further extract biological insights from dynamic networks, DRAFT uses edge clustering on the reconstructed networks to pinpoint shared and stage-specific activity of TFs (regulators) as well as latent Dirichlet allocation (LDA) to assign active gene topics to each stage. Based on functional enrichment of gene topics, we analyze how pathway activities vary over time.

DRAFT fuses prior graph structure and molecular measurements across stages by combining graph attention and recurrent temporal learning (**Fig. 1b**). At each stage, DRAFT uses Graph Attention Networks (GATs^16^) to nonlinearly integrate prior graph topology with gene expression and chromatin accessibility profiles. In each iteration, GAT allows a node to aggregate information from the features of its neighbors and maps nodes into latent-space representations; after multiple rounds of message passing, the features are influenced by multi-hop neighborhood information. A Gated Recurrent Unit (GRU^20^) then propagates information along lineage relationships, linking each node’s current representation to preceding lineage stages (**Fig. 1b**). The GAT and GRU modules are jointly trained to reconstruct prior relationships and produce stage-specific node embeddings (**Methods**). After training, the model scores candidate edges using the cosine similarity in the learned latent space, resulting in a stage-specific regulatory network. Using Gaussian kernel interpolation in the latent space, DRAFT can furthermore reconstruct continuously varying GRNs along the whole time axis (**Fig. 1b; Methods**).

To ease interpretation of the learnt regulatory dependencies, DRAFT performs edge clustering and gene topic analysis (**Fig. 1c**) inspired by prior research^13^. By applying k-means to inferred edges across stages, we identify edge clusters (ECs) and characterize their activity profiles over time, which in turn highlights the TFs associated with temporally important regulatory programs. To identify and trace the most active gene sets over time, DRAFT uses gene topic clustering. We treat each TF–TG edge as a document whose words are the stages at which it is selected, and perform Latent Dirichlet Allocation (LDA) to obtain gene topics and their activity profiles over time, linking them to known pathways and functional annotations.

### DRAFT recovers dynamic GRNs from simulated single-cell multi-omics data with ground truth networks

To evaluate DRAFT’s performance in reconstructing dynamic GRNs, we developed a single-cell multi-omics simulation framework with ground-truth regulatory interactions. Each simulated dataset comprised 50 TFs, 150 TGs and five stages (consecutive segments of one simulated trajectory, T0–T4), with 200 cells per stage. Starting from static prior networks on TF-TG relationships of different topology, we selected stage-specific active edges that change gradually over time. Thereby, the prior network included persistently active edges, edges that were active only at some stages, as well as edges that remained inactive, representing both the lack of temporal specificity and presence of false positives in available regulatory priors. These ground-truth stage-specific TF–TG networks served as the GRN input to scMultiSim^22^ to generate paired gene expression and chromatin accessibility profiles. Each TG was linked to its candidate TFs through synthetic REs **(Methods, Fig. 2a)**.

**Fig. 2.**
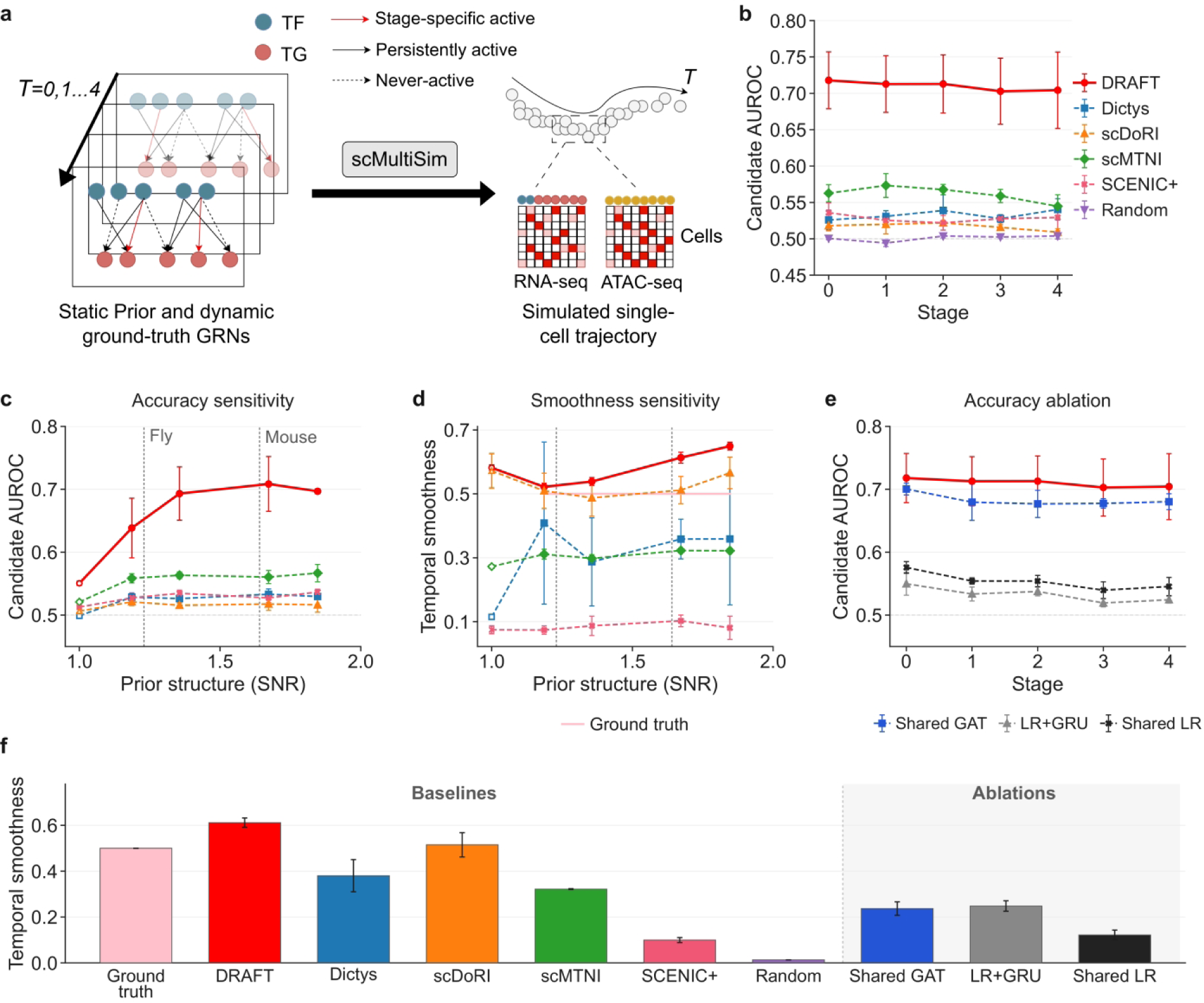
Benchmarking DRAFT on simulated multiome time courses with known GRNs. **a**, Overview of the simulation. Ground-truth GRNs were built for five stages from 50 TFs and 150 target genes (TGs) organized into three modules. In the static prior, each TG has 12 candidate TFs (9 within its module and 3 outside); at each stage 6 of the within-module candidates are active, and 10% of active edges are replaced between adjacent stages, without reversal. Paired single-cell RNA and chromatin accessibility profiles (200 cells per stage) were generated from these GRNs with scMultiSim^22^. Black, red and dashed arrows denote persistently active edges, stage-specific active edges and never-active candidate edges, respectively. **b,** AUROC at each stage for DRAFT, SCENIC+^1^ (TF-to-gene component), scDoRI^12^, scMTNI^13^, Dictys^11^ and a random control, using the ground-truth GRN as the gold standard. All methods received the same observations and priors and were scored on the same candidate TF–TG pairs. **c,d,** Sensitivity of candidate-set AUROC (**c**) and temporal smoothness (**d**) to prior structure, quantified by spectral signal-to-noise ratio (SNR; **Supplementary Methods**). The number of cross-module candidates per TG was varied across 2, 3, 6 and 8, while retaining nine within-module candidates and six active within-module regulators per stage. Vertical dashed lines indicate the SNRs of the fly^23^ and mouse^13^ data priors. Temporal smoothness is 1 − the mean Frobenius distance between networks at adjacent stages divided by the largest pairwise distance, on the candidate pairs **(Methods)**. Open symbols denote the independent random-network experiment from **Supplementary** Fig. 2, plotted at SNR = 1 only as a reference**. e,** AUROC comparison between DRAFT and three internal ablations: Shared GAT, which removes the GRU; LR+GRU, which replaces the GAT with linear encoders; and Shared LR, which combines both changes. **f,** Temporal smoothness of DRAFT, the baseline methods, the random control and ablations; ground truth is shown as a reference. The shaded region identifies the internal ablations. Results use 3 independently simulated datasets, with 5 model seeds per dataset. Values are means across dataset-level averages, and error bars indicate the standard deviation across the three datasets. Panels **b,e,f** show the main design (SNR = 1.67 in **c,d**).

We benchmarked DRAFT against scDoRI^12^, scMTNI^13^, SCENIC+^1^, Dictys^11^ and a random control **(Methods)**. DRAFT and all baseline methods received the same static prior network and measurements at every stage. Predictive performance was quantified by the area under the receiver operating characteristic curve (AUROC) over all candidate TF–TG pairs of the prior network, with currently active edges treated as positives and inactive edges as negatives. Network temporal smoothness was defined based on the Frobenius distance between consecutive networks **(Supplementary Fig. 1)**. To assess the contributions of individual model components, we evaluated three ablated variants of DRAFT, removing the graph attention network, the gated recurrent unit or both (**Methods**).

Across different simulation parameters, we observed a clear gain in accuracy and smoothness of the inferred networks. This gain was most pronounced for highly structured prior networks. While on randomly generated regulatory networks, DRAFT achieved a mean AUROC of 0.550, only slightly higher than ablated versions (Shared GAT 0.534; LR+GRU 0.531; Shared LR 0.531) or other methods (AUROCs ranging from 0.499 to 0.521) **(Supplementary Fig. 2),** with modular priors, the gain was larger **(Fig. 2b,c)**. Here, DRAFT achieved a mean AUROC of 0.710, exceeding scMTNI (0.561), Dictys (0.533), SCENIC+ (0.528) and scDoRI (0.517; **Fig. 2b**). DRAFT also attained higher mean network smoothness (0.612) than scMTNI, Dictys, SCENIC+ and scDoRI, which scored 0.322, 0.380, 0.100 and 0.515, respectively **(Fig. 2f)**. This dependence on prior structure is consistent with DRAFT’s use of graph attention to aggregate neighborhood information, beyond constraining or weighting candidate edges like baseline methods. Ablation analysis supported complementary contributions of the two modules: graph attention was associated primarily with improved candidate-edge ranking, whereas recurrent learning contributed more strongly to temporal smoothness and provided a modest additional gain in AUROC **(Fig. 2e,f)**. Sensitivity to the edge-turnover rate and the fraction of inactive candidates is shown in **Supplementary Fig. 3**.

Finally, we extended the simulation to a trajectory with a shared trunk with two daughter branches, propagating GRU states along parent–child relationships while retaining the prior construction and training objective. DRAFT achieved average AUROC 0.679 and smoothness 0.652, exceeding scMTNI and all three ablations on both mean metrics **(Methods; Supplementary Fig. 4)**.

### DRAFT reveals branch-specific regulatory programs during mouse cellular reprogramming with unpaired multi-omics data

To further evaluate DRAFT’s ability to characterize regulatory dynamics during cell-state transitions, we applied it to an unpaired dataset of mouse embryonic fibroblast (MEF) reprogramming, comprising separate scRNA-seq^24^ and scATAC-seq profiles^13^. Four intermediate sampling times (days 3, 6, 9 and 12), together with MEF and mESC endpoints, gave six sampling times. Following the scMTNI preprocessing pipeline^13^, we obtained seven cell clusters forming successful (C4–C8–C1–C5) and unsuccessful (C4– C8–C7–C3–C2) trajectories from the MEF-enriched starting cluster C4 **(Fig. 3a)**, together with TF–TG priors derived from motif binding score and genomic distance information **(Methods)**.

**Fig. 3.**
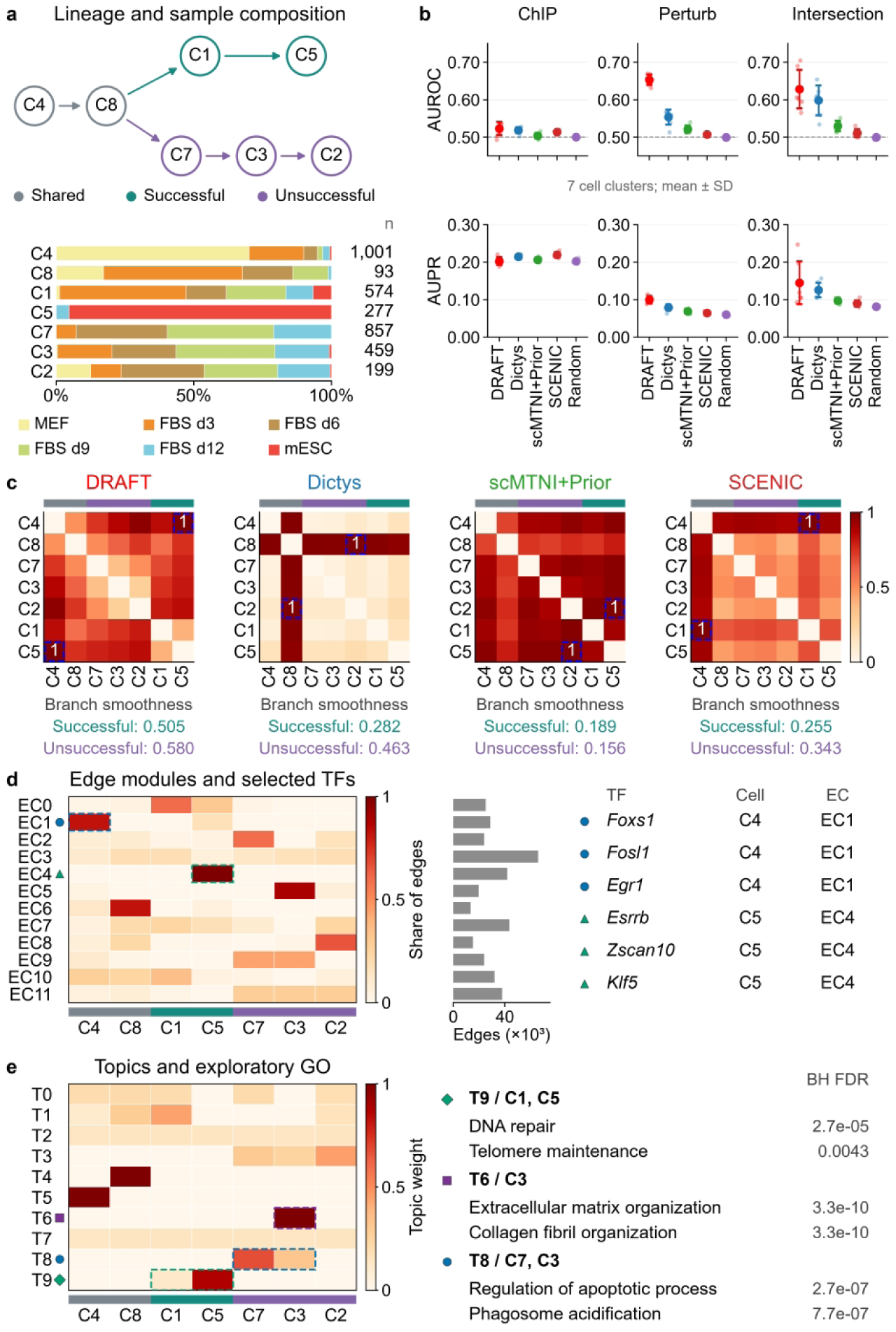
Cell-cluster–specific GRN inference with DRAFT from mouse cellular reprogramming data. **a**, The cell lineage structure together with, for each cell cluster, the proportion of cells contributed by each sample^13^. Arrows indicate lineage relationships connecting shared clusters C4/C8 to the successful branch (C1→C5) and unsuccessful branch (C7→C3→C2). Grey, teal and purple denote shared, successful and unsuccessful lineage segments, respectively. Stacked bars show the proportion of cells contributed by each sample (MEF, FBS days 3, 6, 9 and 12, and mESC) within each cell cluster. Numbers at right indicate cell counts. **b,** DRAFT, Dictys, scMTNI+Prior, SCENIC and Random were evaluated against mESC ChIP-seq, knockdown perturbation and their intersection gold standards adopted from the original study^13^. Rows show AUROC (top) and AUPR (bottom). Small points represent seven cell clusters; large points and error bars show the mean ± sample SD. scDoRI was not included because it only accepts paired data as input. **c,** Pairwise Frobenius distances between cluster-specific regulatory networks inferred by DRAFT, Dictys, scMTNI+Prior and SCENIC, normalized by each method’s maximum pairwise distance across the seven cell clusters. Blue dashed boxes mark the maximum-distance pair and its symmetric entry, labeled 1. Below each heatmap, branch smoothness along the successful and unsuccessful lineages (defined as in Fig. 2, with distances restricted to each branch). **d,** Distribution of 12 k-means-derived edge clusters (ECs) across cell clusters, normalized within each EC. Bars show total edge occurrences across cell clusters. Selected candidate transcription factors (TFs) with high network centrality are listed with their associated cell clusters and dominant ECs. **e,** Row-normalized weights of 10 LDA-derived gene topics across cell clusters, alongside selected enriched Gene Ontology biological process terms. BH FDR denotes Benjamini–Hochberg-adjusted *P* values. Dashed boxes and matching markers in **d,e** link selected heatmap entries to TF and functional annotations.

We compared DRAFT with scMTNI, Dictys, SCENIC and a random control against ChIP-based interactions from ESCAPE and ENCODE, perturbation-based interactions combining ESCAPE loss/gain-of-function data with Nishiyama et al., and their intersection^25–27^ following the scMTNI benchmark^13^ (**Methods**). Performance differences among methods were modest on the ChIP benchmark. On the perturbation benchmark, which provides evidence of functional regulatory effects, DRAFT achieved higher AUROC (0.653) and AUPR (0.100) than all baselines. DRAFT also achieved the highest mean AUROC (0.628) and AUPR (0.145) on the ChIP–perturbation intersection, although its advantage over Dictys was not statistically significant (**Fig. 3b**). In pairwise network comparisons, DRAFT showed the highest smoothness along both branches and predicted the MEF-enriched C4 and mESC-enriched C5 networks as the most divergent pair, consistent with their distinct cell states^13,28^, whereas maximum-distance pairs differed among other methods (**Fig. 3c**).

To further dissect regulatory differences between successful and unsuccessful reprogramming, we performed edge clustering and LDA-based gene-topic discovery, inspired by scMTNI^13^ **(Methods, Fig. 1c)**. Edge clusters (ECs) group edges by their distributions across cell clusters, allowing the same TF to occur in multiple ECs through distinct subsets of inferred targets. Within each EC, we ranked TFs by centrality, defined as the number of unique target genes linked to each TF (**Methods**). Edge clustering highlighted a starting-state module, EC1, containing *Foxs1*, *Fosl1* and *Egr1*, and a terminal-state module, EC4, containing *Esrrb*, *Klf5* and *Zscan10* (**Fig. 3d**). These associations agree with reported roles of these TFs: repression of *Fosl1* contributes to somatic-enhancer silencing, whereas its overexpression impedes reprogramming^28^. *Foxs1* is implicated in myofibroblast responses^29^. *Egr1* knockdown can enhance fibroblast reprogramming^30^. In contrast, *Klf5* supports mouse ESC self-renewal and reprogramming toward naïve pluripotency^31^, while *Esrrb* promotes ESC self-renewal and the acquisition of pluripotency^32^. *Zscan10* (also known as *Zfp206*) participates in the Oct4–Sox2 transcriptional regulatory network in ESCs^33^. In addition, EC3 and EC7 were broadly distributed across reprogramming states and contained *Sp1*, *Sp3* and *Gabpa* **(Supplementary Fig. 5)**. Previous studies linked *Sp1* to cell-cycle-associated enhancement of reprogramming^34^, *Sp3* to repression of cardiomyocyte-specific genes in cardiac fibroblasts^35^ and *Gabpa* to the establishment of naïve pluripotency during mouse embryonic development^36^.

Using gene topic discovery, we identified ten gene topics and examined their cluster distributions and functional enrichment (**Fig. 3e**, **Supplementary Fig. 6**). T9 was predominantly associated with the mESC-enriched terminal cluster C5 and enriched for DNA repair and telomere maintenance (BH FDR = 2.7 × 10^−5^ and 4.3 × 10^−3^, respectively), consistent with established links between genome maintenance and cellular reprogramming^37,38^. In contrast, T6 was concentrated in C3 on the unsuccessful branch and enriched for extracellular matrix organization and collagen fibril organization (BH FDR = 3.3 × 10^−10^ for both), consistent with retention of fibroblast- or mesenchymal-associated matrix programs^39,40^. T8 was also associated with the unsuccessful branch (C7/C3) and enriched for regulation of the apoptotic process and phagosome acidification (BH FDR = 2.7 × 10^−7^ and 7.7 × 10^−7^, respectively). The latter annotation comprised ten V-ATPase subunit genes; these enrichments suggest a candidate cell-survival and intracellular-acidification signature without establishing whether these processes were activated or suppressed. Cell-cycle terms were enriched in T7, whose broad distribution across clusters indicated that this program was not exclusive to successful reprogramming.

### DRAFT reconstructs dynamic eGRNs and early stem-cell regulatory programs during spermatogenesis in the adult *Drosophila* testis from paired multiome data

The *Drosophila* testis, with its well-defined anatomy and a temporally continuous progression of stages, provides an ideal testbed for DRAFT. At its apex, germline stem cells (GSCs) and cyst stem cells (CySCs) surround hub cells to form a niche regulated by JAK/STAT^41^, bone morphogenetic protein (BMP)^42,43^ and Hh^23^ signaling. Each gonialblast (Gb) is then enclosed by two non-dividing cyst cells, which grow and mature in lockstep with the germ cells and support them through four transit-amplifying divisions, meiosis and spermiogenesis^44^. We used published single-nucleus multiome data (paired snRNA-seq and snATAC-seq) for 10,335 annotated nuclei from the apical end of the adult testis^23^. Following the original study, we performed clustering and resolved the germline and somatic lineages and rare niche hub cells. Stage labels follow the original annotation, which partitioned germline and somatic latent time into equal-width bins. The germline spans GSCs, Gbs, 2–16-cell spermatogonia (Sp-2/4/8/16), and primary and secondary spermatocytes (1° Sc and 2° Sc). Because secondary spermatocytes are transient whereas primary spermatocytes grow for ∼3.5 days, the six “2° Sc” bins, which contain the most nuclei, may represent late primary spermatocytes progressing towards and through meiosis **(Fig. 4a)**. The somatic lineage spans CySCs and early/late cyst cells (eSCC/lSCC), before branching into two differentiation branches, SdA/SdB **(Fig. 4a)**. We constructed regulatory priors per lineage from the top 3% of TF–RE pairs ranked by cisTarget motif-enrichment scores^1^ and genomic proximity-based RE–TG pairs (**Methods**).

**Fig. 4.**
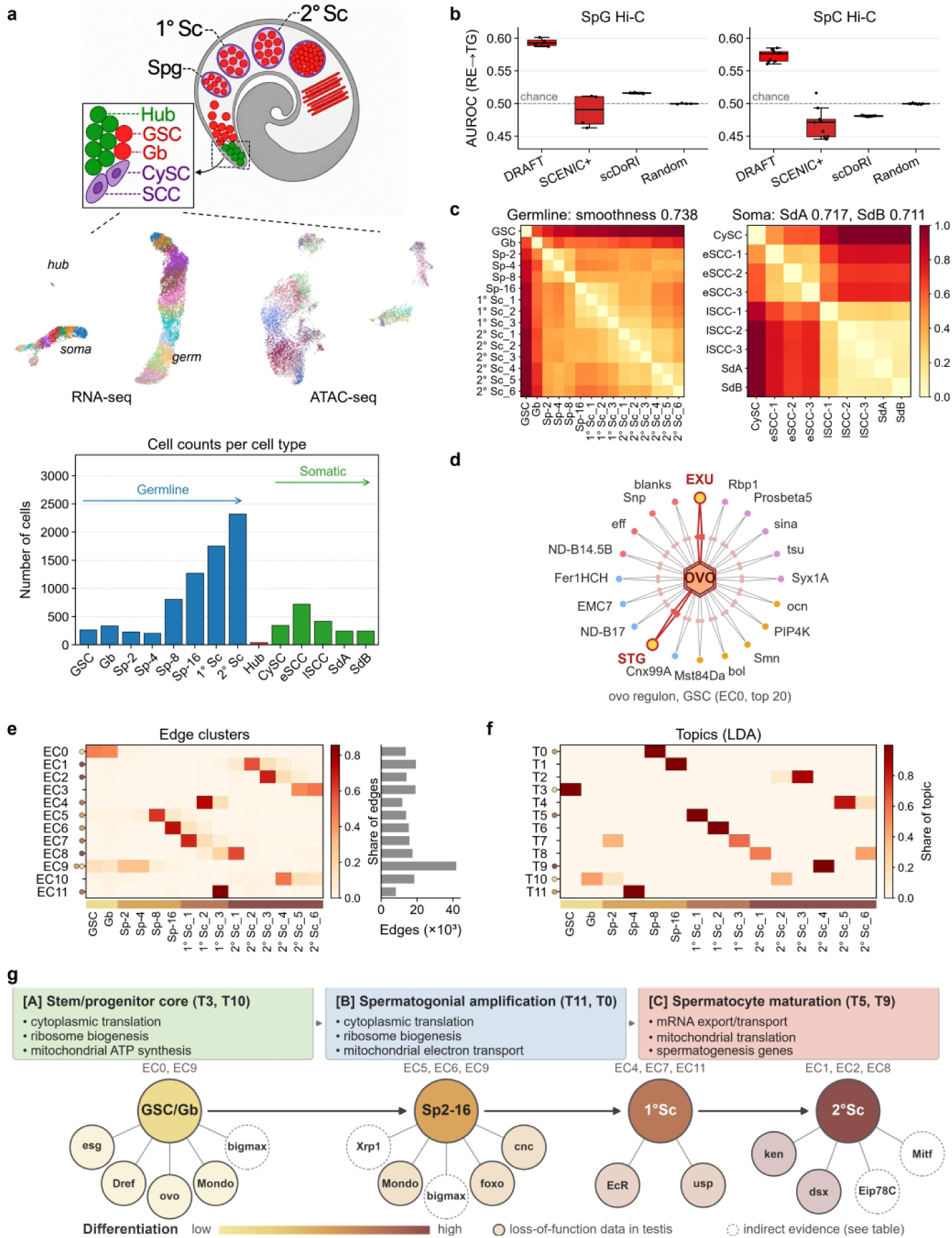
Application of DRAFT to paired *Drosophila* adult testis data. **a**, Overview of the *Drosophila* testis dataset^23^. Top, testis schematic with the germline and somatic lineages; inset, niche and early progenitors. Middle, UMAPs of snRNA-seq (left) and snATAC-seq (right) cells, colored by cluster; germline and somatic cells form separate trajectories. Bottom, cells per cell type. Stage labels as in ref. 23. **b,** Hi-C benchmark at the spermatogonial (SpG, left) and spermatocyte (SpC, right) stages: AUROC for ranking Hi-C-supported RE–TG pairs above an equal number of other pairs, all within the shared RE–TG prior. Each point is one stage (SpG, Sp-2 to Sp-16, n = 4; SpC, 1° Sc_1 to 2° Sc_6, n = 9). Boxes show the median and interquartile range; whiskers extend to the most extreme values within 1.5× the interquartile range. Dashed line, chance. SCENIC+ region-to-gene models were fitted per stage (Methods). scMTNI and Dictys (TF–TG networks only) are not shown. **c,** Frobenius distance between DRAFT’s TF–TG networks at each pair of stages, divided by the largest distance. Smoothness (titles) is as defined in Fig. 2; for the soma, along each branch (CySC to lSCC-3, then SdA or SdB). **d,** GSC-stage eRegulon of *ovo*: the top 20 targets (CG-numbered genes omitted) in its edge cluster EC0, each linked through its two top-ranked REs. Red, targets with literature support. **e,** Activity profile of the edge clusters across stages (k-means, 12 clusters): heatmap, fraction of each cluster’s edges present at each stage; bars, edges per cluster (counted at every stage). The strip below groups the stages as the nodes in **g**; dots mark the clusters whose most central TFs appear in **g**. **f,** Activity of the LDA topics (12) across stages: each topic’s weight over stages, normalized to sum to one; strip and dots as in e (dots, topics summarized in the boxes of g). **g,** Summary of the germline trajectory. Nodes are stage groups, darker with differentiation, labeled with their edge clusters; GSC and Gb are merged because they share EC0. Badges, TFs among the ten most central in these clusters (**Supplementary** Fig. 8): solid, loss-of-function data in the Drosophila testis (published or this study); dashed, indirect evidence only (binding partner, ovary data or activity reporter; **Supplementary Table 2**). Boxes, GO terms enriched in the listed topics (**Supplementary** Fig. 9).

We reconstructed stage-specific eGRNs with DRAFT, scDoRI, SCENIC+ and a random control **(Methods)**. To assess network quality, we used independent Hi-C maps from adult *bam*Δ86 mutant testes enriched in spermatogonia-like germ cells and from wild-type third-instar larval testes enriched in primary spermatocytes^45^. TSS–region contacts (50,683 and 44,748, respectively) served as a permissive reference for spermatogonial and spermatocyte stages. Because chromatin contacts indicate proximity rather than regulation and both samples contain somatic cells, AUROC values are expected to be modest. Within the RE–TG prior shared by all methods, DRAFT ranked Hi-C-supported pairs above randomly sampled prior pairs with mean AUROCs of 0.59 and 0.57 across spermatogonial and spermatocyte stages, compared with 0.49 and 0.47 for SCENIC+, 0.52 and 0.48 for scDoRI, and 0.50 for the random control (**Fig. 4b**). Regarding temporal smoothness, DRAFT produced gradually changing networks, with smoothness scores of 0.74 for the germline and 0.72 and 0.71 for the SdA and SdB somatic branches **(Fig. 4c)**. These scores were higher than those of SCENIC+ (0.67, 0.53 and 0.53) and random networks (0.00), whereas scDoRI scored highest (0.90, 0.80 and 0.81) (**Supplementary Fig. 7**). However, each scDoRI stage network mixes fixed topic networks by stage-specific proportions; with random topic networks and the same proportions, smoothness still reached 0.85, 0.75 and 0.75, indicating that scDoRI’s smoothness largely reflects the topic proportions rather than the network edges.

Edge clustering of the stage-specific networks **(Methods)** yielded 12 edge clusters (EC0–EC11) with clear temporal preferences, from GSC/Gb and early spermatogonia to late spermatogonia (Sp-8/Sp-16) and spermatocytes **(Fig. 4e)**. Within each cluster, we ranked TFs by centrality **(Supplementary Fig. 8)**.

We summarized each developmental window by the top-10 TFs of its edge clusters that have functional support **(Fig. 4g and Supplementary Table 2)**. In GSC/Gb-associated EC0 and EC9, DRAFT recovered two established regulators: *Dref*, required for male GSC maintenance^46^, and *ovo*, validated by CRISPR knockout as a GSC regulator^23^ and required for germline development in both sexes^47^. It also recovered *esg*, which is expressed in male GSCs^48^ and required in hub cells^49^ and CySCs^50^ for niche maintenance but not intrinsically in GSCs^49^. *Mondo* led EC9 (GSC–Sp-4), with its binding partner *bigmax* (*Drosophila Mlx*) ranking fifth^51^. During spermatogonial amplification (Sp-2 to Sp-16; EC5, EC6 and EC9), *Mondo* and *bigmax* were joined by *foxo* and *cnc*: *foxo* mutant testes accumulate undifferentiated germ cells^52^, and knockdown of *cnc*, the *Nrf2* orthologue, in spermatogonia alters transit-amplifying divisions in an oxidative-stress pathway^53^. The most central TF in EC5 (Sp-8), *Xrp1*, has to our knowledge not been studied in the testis, but suppresses translation via eIF2α phosphorylation and participates in cell competition and responses to DNA damage in other tissues^54,55^. Together with the Sp-4 translation/ribosome-biogenesis topic (T11), these candidates suggest nutrient-sensing and redox-related programs during amplification, with a possible Xrp1-linked fitness role requiring validation. The original SCENIC+ analysis^23^ also highlighted *ovo, esg* and *foxo* **(Supplementary Table 3)**.

In spermatocytes, the top-ranked TFs shifted to nuclear receptors and sex-determination regulators **(Fig. 4g)**. *EcR* ranked first in EC4 (1° Sc), alongside its partner *usp*; *ken* and *dsx* ranked first in 2° Sc-associated clusters (EC1, EC2 and EC8), followed by the ecdysone-induced nuclear receptor *Eip78C* and the lysosomal regulator *Mitf*. In the testis, loss-of-function data for *EcR, usp, ken* and *dsx* come mainly from somatic cells: ecdysone signalling through *EcR* and *usp* maintains CySCs and, non-autonomously, GSCs^56^. *ken* is required for CySC self-renewal^57^, and *DsxM* maintains the CySC male identity^58^. For the remaining TFs, evidence of a germline role is indirect: *Eip78C* is required in the female germline^59^, and *Mitf* activity rises in meiotic spermatocytes^60^. *EcR, usp* and *Eip78C* outline an ecdysone-response axis, but these TFs, which SCENIC+ also highlighted **(Supplementary Table 3)**, remain network-inferred candidates rather than established spermatocyte regulators.

LDA topic modeling yielded 12 gene topics (T0–T11; **Fig. 4f**), ten with significant GO terms that broadly match known stage features **(Supplementary Fig. 9)**. GSC and Gb topics (T3 and T10) were dominated by cytoplasmic translation, ribosome biogenesis and mitochondrial ATP synthesis, consistent with the roles of ribosome biogenesis in GSC differentiation^61^ and of mitochondrial function in male GSC maintenance^62^. In addition, the GSC topic was enriched for male germline stem cell population maintenance. Translation and ribosome biogenesis remained prominent in the spermatogonial topics (T11: Sp-4; T0: Sp-8), consistent with the high biosynthetic demand of transit-amplifying divisions. Spermatocyte topics shifted towards RNA processing, with mRNA export and mitochondrial translation in T5 (1° Sc) and mRNA splicing in T6 (1° Sc), consistent with the extensive transcription and mRNA processing that prepare spermatocytes for meiosis and spermatid differentiation^63^. T9 (2° Sc_4) was enriched for spermatogenesis and male gamete generation **(Fig. 4g)**.

We further decomposed the networks into eRegulons, using *ovo* (rank 9 in EC0; **Supplementary Fig. 8**) as an example. Its transcripts are restricted to GSCs and Gbs and decline as *bag of marbles* (*bam*) transcripts rise in the adult testis^23^. The GSC-stage ovo eRegulon **(Fig. 4d)** contained *stg*, an Ovo target validated by smFISH in the testis^23^, and *exu*, a gene bound by Ovo in the female germline^64^.

### DRAFT predicts an early germline role for Mondo that precedes its expression peak and is supported by germline knockdown

Comparing TFs highlighted by DRAFT with those identified in the original study revealed regulators whose network-centrality peaks differed from their RNA-expression peaks **(Supplementary Table 4)**. We hypothesized that combining graph attention with temporal memory could help reveal candidate regulatory roles that are not readily apparent from RNA-expression timing alone. In particular, *Mondo* maintained a high predicted GRN centrality throughout the early germline lineage (GSC–Sp-4), suggesting a candidate early germline role. However, its gene expression peaked at a later developmental stage (1° Sc) **(Fig. 5a, Supplementary Fig. 8)**. To investigate the temporal mismatch, we performed two ablations: removing GRU and replacing GAT with linear encoders while retaining GRU (**Supplementary Fig. 10**). Centrality in SCENIC+-inferred networks per stage served as an external reference. Both GAT-containing models recovered *Mondo* network connectivity at the GSC stage, whereas other models were zero at this stage. Meanwhile, the predominant centrality peaks of GRU-containing models occurred early, whereas GRU removal shifted the predominant peak later, closer to the RNA peak. These comparisons suggest complementary contributions: graph encoding may support early *Mondo* connectivity, whereas temporal recurrence may concentrate *Mondo*’s predicted role within an early window.

**Fig. 5.**
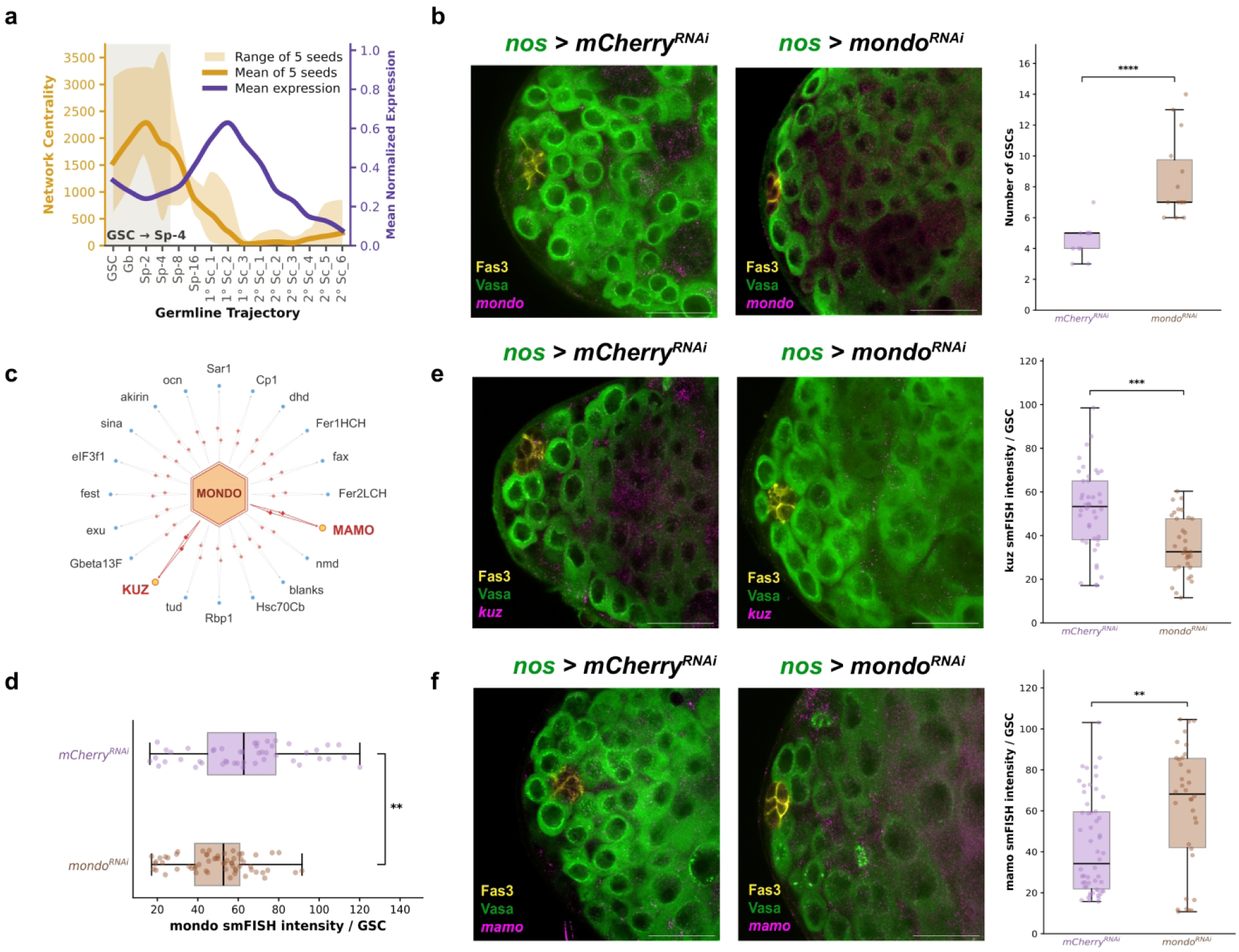
DRAFT predicts a candidate early germline role for *Mondo* that is supported by germline knockdown. **a**, Orange: network centrality of Mondo; the band is the range and the thick line the mean of five models trained with five different random seeds. Purple: mean normalized expression of *Mondo*. In all seeds, the stage of maximal centrality lies between GSC and Sp-4 (grey) and precedes the RNA expression peak. **b**, Representative images of the testis apex in *nos-GAL4::VP16 > UAS-mCherry RNAi* (control) and *nos-GAL4::VP16 > UAS-Mondo RNAi* males stained for Fas3 (hub, yellow) and Vasa (germ cells, green), with *Mondo* smFISH (magenta). Right, number of GSCs per testis, scored by 3D segmentation as Vasa-positive germ cells in contact with the Fas3-positive hub **(Methods)**. Each dot represents one testis; n = 11 (control) and 14 (Mondo RNAi) testes from 2 independent experiments. Two-sided Mann-Whitney U test, P = 8.30 × 10^−5^. **c**, Representative early germline Mondo eRegulon (GSC–Sp-4). Target-gene ranks were combined across the continuous networks from GSC to Sp-4 by the geometric mean, and the 20 most consistently ranked targets are shown with their associated regulatory elements (small symbols on edges). *kuz* and *mamo* are highlighted. **d**, *Mondo* smFISH intensity per GSC in control and Mondo RNAi testes (n = 47 GSCs from 9 control testes and n = 61 GSCs from 9 Mondo RNAi testes). Two-sided Mann-Whitney U test comparing individual GSC measurements, P = 0.003805. **e, f**, Representative images and quantification of *kuz* (**e**) and *mamo* (**f**) smFISH (magenta) with Fas3 (yellow) and Vasa (green) in control and Mondo RNAi testes. Plots show smFISH intensity per GSC; for *kuz* (**e**), n = 42 GSCs from 8 control testes and n = 33 GSCs from 5 Mondo RNAi testes. Two-sided Mann–Whitney U test comparing individual GSC measurements, P = 0.000109. For *mamo* (**f**), n = 54 GSCs from 5 control testes and n = 34 GSCs from 10 Mondo RNAi testes. Two-sided Mann–Whitney U test comparing individual GSC measurements, P = 0.006345. Box plots show the median and interquartile range; whiskers extend to the extreme data points within 1.5 interquartile range (IQR). Scale bars, 20 µm. *P < 0.05, **P < 0.01, ***P < 0.001, ****P < 0.0001.

To assess the biological relevance of the early role predicted by DRAFT, we knocked down *Mondo* in the germline and measured the expression of predicted downstream genes. *Mondo* is a conserved, sugar-activated TF that acts together with its partner Mlx (encoded by *bigmax*) and controls most sugar-responsive genes in *Drosophila*, including the glycolytic regulator PFK2^51,65^. Nutrient signalling is known to influence GSC behaviour: protein starvation reduces GSC numbers in the testis^66,67^, and a high-sugar diet reduces GSC proliferation in the ovary^68^. Germline-specific knockdown of *Mondo* (*nos-GAL4::VP16 > UAS-Mondo^RNAi^*) increased the number of germ cells contacting the hub compared with the *nos-GAL4::VP16 > UAS-mCherry^RNAi^* controls **(Fig. 5b)**. Because our scoring is positional, this increase could reflect either an expanded GSC pool or an accumulation of GSC-like cells that fail to initiate differentiation, as reported after germline knockdown of mTor^69^. In addition, because knockdown was not restricted to adulthood, an effect arising during gonad development cannot be excluded. In either case, the phenotype supports a candidate role for *Mondo* in the early germline.

To define the early germline *Mondo* eRegulon, we integrated TG rankings across the continuous regulatory networks from GSC to Sp-4 using the geometric mean and selected the top 20 most consistently ranked downstream targets **(Fig. 5c)**. Among them, *kuzbanian (kuz)* and *mamo* were prominent candidates **(Supplementary Table 5)**. *kuz* encodes Kuzbanian, the ADAM10-family metalloprotease that performs the S2 cleavage of Notch in the signal-receiving cell and also processes the ligand Delta^70^. In the testis, Delta from the germline activates Notch in the somatic cyst cells of the transition zone, and this signalling is required for survival of the germline lineage^71^. Because germ cells are the ligand-sending cells in this interaction, a germline role for Kuz would most plausibly involve Delta processing or a Notch-independent substrate, which remains to be tested. *mamo* encodes a BTB/POZ zinc-finger protein. A truncated version of Mamo activates *vasa* expression in synergy with OvoB in the primordial germ cells^72^. Germline *Mondo* knockdown reduced the *Mondo* smFISH signal in GSCs by 22% **(Fig. 5d)**, confirming knockdown in the targeted cells, and decreased *kuz* and increased *mamo* transcript levels **(Fig. 5e,f)**.

*Mondo* is not the only TF with this pattern. Applying the two criteria used to select *Mondo*, a temporal offset between network centrality and mean RNA expression and a rank among the ten most central TFs of its edge cluster, identified 22 further TFs **(Supplementary Table 4)**. Temporal offsets alone do not establish biological relevance, because randomly re-paired profiles produced at least as many offsets **(Supplementary Fig. 11)**. Offset directions were largely reproducible across independent initialisations **(Supplementary Fig. 11; Supplementary Table 6)**. Architectural ablations and randomization of TF degrees in the prior both reduced agreement with the original peak stages **(Supplementary Fig. 11)**. We therefore treat these 22 TFs as candidates. A literature search identified stage-resolved functional evidence for two of them, and in both cases it agrees with the network-inferred timing rather than with the expression peak. *Dref* centrality peaks in GSCs, although its mRNA peaks in spermatocytes and SCENIC+ assigns it to spermatocytes **(Supplementary Table 3)**. Intriguingly, *Dref* has been shown to be required for GSC self-renewal in the male germline^46^. Conversely, *Mitf* centrality peaks in spermatocytes although its mRNA peaks in GSCs, and a *Mitf* activity reporter is indeed higher in spermatocytes than in spermatogonia^60^. These findings illustrate how dynamic GRN inference can point to regulatory activity that is not apparent from expression timing and help direct experiments.

### Aligning germline and somatic GRNs in continuous time highlights stage-specific ligand–receptor communication

Finally, we hypothesized that the dynamic GRNs inferred by DRAFT could further allow predictions for time-resolved cell-cell communication (CCC) across lineages. In testis development, hub cells as quiescent niche cells^73^ can be seen as constant senders while somatic and germline cells develop together^73,74^ and are in direct contact at corresponding developmental stages. Previous analyses of this dataset identified cell-type-level ligand–receptor (LR) interactions^23^ but without resolving their developmental timing. To analyze time-varying signaling, somatic-cell stages must be matched to those of the enclosed germ cells. However, the two pseudotime trajectories were inferred separately, and developmental progression may not be synchronized or uniform, preventing direct stage correspondence. Existing single-cell trajectory alignment methods mainly use gene expression to align the same biological process across datasets or conditions and are not designed for distinct lineages co-developing within one tissue^75,76^. We therefore used dynamic time warping (DTW)^77^ based on the two lineages’ dynamic networks inferred by DRAFT to nonlinearly map the somatic onto the germline trajectory, enabling communication analysis along a common time axis **(Fig. 6a, Supplementary Table 7)**. The alignment path was fixed at both endpoints and subjected to monotonic smoothing **(Methods)**.

**Fig. 6.**
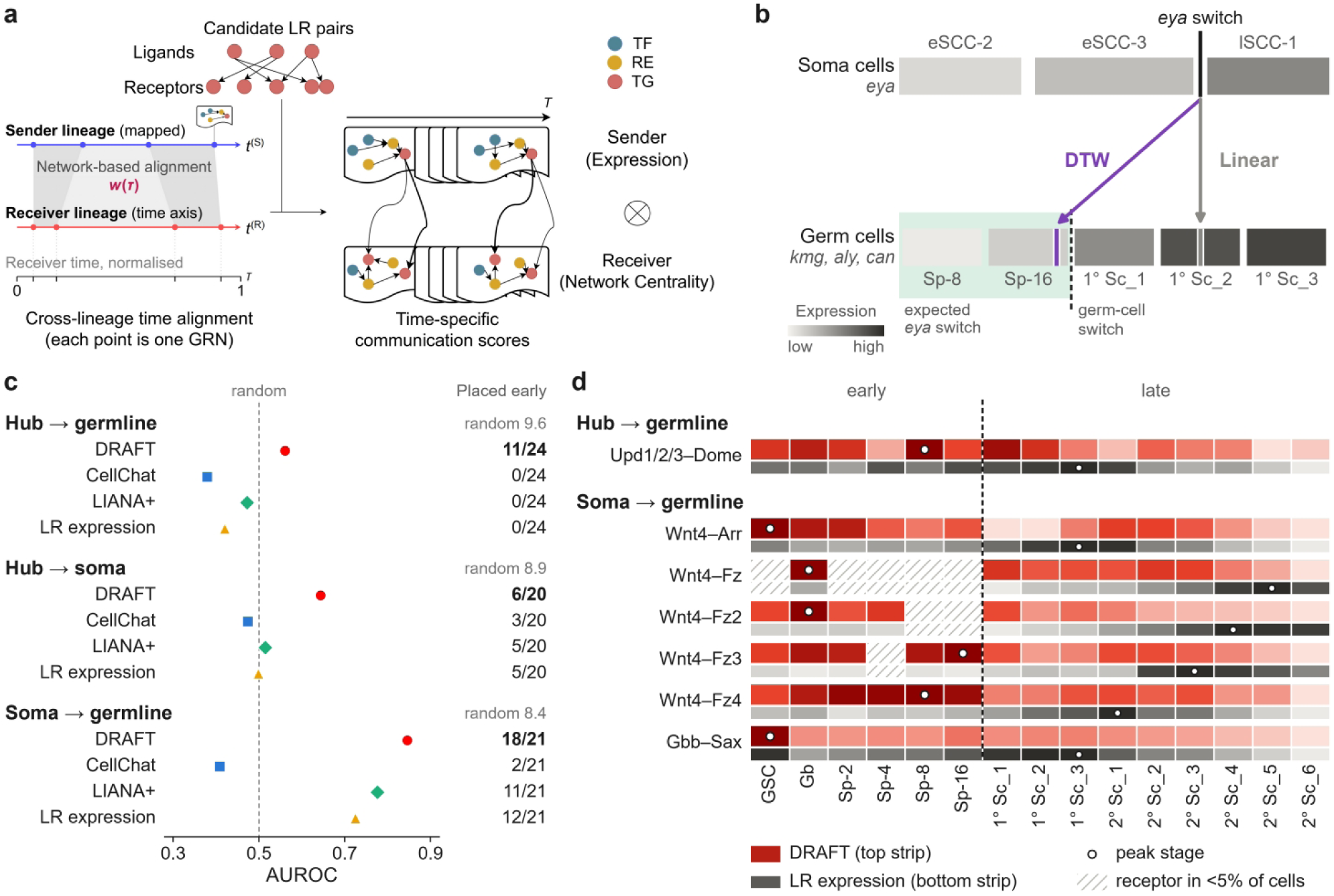
Inter-lineage time alignment and stage-specific cell-cell communication prediction. **a**, Time alignment and LR scoring. Sender stages are mapped onto the receiver’s time axis (τ) by dynamic time warping. At each receiver stage, every candidate LR pair is scored as ligand expression in the sender multiplied by the receptor’s centrality in the receiver’s stage-specific network. Arrow width indicates the LR score, which changes along τ. **b,** Alignment check with an anatomical anchor. Top, *eya* in cyst cells; bottom, mean of spermatocyte-program genes (*kmg*, *aly*, *can*) in germ cells. Black bar and dashed line, the switch in each lineage (largest increase between adjacent stages). Arrows, the *eya* switch mapped onto germline stages by DTW (purple) and by linear alignment (grey). Green, expected window, as Eya rises in cyst cells from late spermatogonia. **c,** AUROC of the earliness score **(Methods)** for literature-supported LR pairs versus other candidate pairs; dashed line, random. Labels, literature pairs peaking in early stages; “random”, number expected by chance. DRAFT germline receivers, the germline model used throughout. **d,** Literature pairs placed early by DRAFT and late by the LR expression product in all five germline models. Top strip, DRAFT; bottom strip, LR expression; each strip is min–max scaled across stages; it shows where the score peaks, not how much it varies. Circles, peaks; hatched, receptor in <5% of cells; dashed line, early/late boundary.

To evaluate the resulting alignment in the absence of ground-truth correspondences, we used a known developmental expression switch and compared DTW with linear alignment, which assumes constant developmental rates and shared start and end times. Eya expression in cyst cells increases as germ cells transition from spermatogonia to spermatocytes and is required for spermatocyte development^78,79^. Its switch is therefore expected between late spermatogonia (Sp-8) and the onset of the germline spermatocyte program. A biologically plausible alignment should map the *eya* switch to this window. We defined both switches by the largest adjacent-stage expression increase. The *eya* switch in cyst cells was located between eSCC-3 and lSCC-1, whereas the switch of the spermatocyte program (*kmg*^80^, *aly*^81^, and *can*^82^) in germ cells was located between Sp-16 and 1° Sc_1 (observed in 94% and 83% of 200 cell-resampling iterations, respectively). DTW mapped the *eya* switch to 0.2 stages before the germline switch, placing it within the expected window. In contrast, linear alignment placed it 1.5 stages after the germline switch (1° Sc_2) **(Fig. 6b)**. Thus, at this transition, DTW provided a correspondence that was more consistent with known biology than linear alignment.

Using the unified time axis, we scored candidate LR pairs from FlyPhoneDB^83^ at each receiver stage as mean sender ligand expression multiplied by receptor network centrality in the receiver’s stage-specific network **(Fig. 6a, Methods)**. These scores are correlative predictions and do not establish causal signaling interactions. Hub ligand expression was constant, whereas ligand expression in somatic sender cells was taken from the aligned sender position. For evaluation, we curated literature-supported signaling pairs from published studies of adult testis signaling pathways^23^ (**Supplementary Table 8**): 24 hub-to-germline pairs, 20 hub-to-soma pairs, and 21 soma-to-germline pairs, with reported functions in GSCs or CySCs. Pairs were classified as early if their scores peaked at GSC– Sp-16 in germline or CySC–eSCC-3 in soma. AUROC assessed discrimination of these reference pairs from other candidates. As controls, CellChat^84^, LIANA+^85^, and the ligand– receptor expression product were used to score the same candidate pairs under identical stage groupings, expression gating, and alignment. Hub-related directions required no alignment and therefore assessed receptor scoring alone, whereas soma-to-germline scoring combined alignment with receptor scoring.

Interestingly, CellChat and LIANA+ showed 93–96% agreement with the expression product in early-versus-late classification, whereas DRAFT showed 73%, indicating that receptor network centrality provided temporal information distinct from expression levels **(Supplementary Fig. 12)**. Whether these differences better reflected known biology was evaluated using the literature-supported pairs. In the hub-to-germline direction, expression-based methods assigned 0/24 reference pairs early (AUROC 0.38-0.47), placing receptors late because their expression increased with the global transcriptional rise in spermatocytes^63^. DRAFT receptor centralities did not show this increase and assigned 11/24 pairs early (average AUROC 0.60; **Fig. 6c, Supplementary Fig. 13**). However, the number of early pairs remained within the range expected by chance (95% range: 5-14), indicating a limited advantage. For hub-to-soma, DRAFT achieved AUROC 0.64 versus 0.47–0.51 for other methods. Because hub cells directly contact only GSCs and CySCs^73^ and these scores lacked spatial information, late hub-derived signaling scores reflected only the signaling reception potential of the receiver cells. In the aligned soma-to-germline analysis, DRAFT assigned 18/21 pairs early (AUROC 0.85), outperforming LIANA+ (11/21; AUROC 0.78), the expression product (12/21; 0.72) and CellChat (2/21; 0.41). Germline-to-soma evaluation was limited to Spi–Egfr, preventing AUROC calculation **(Supplementary Table 8)**. The expression product favored early timing and DRAFT late timing, but early/late maxima differed by only 10.5% and 5.9% of their respective peak scores, so neither timing call is decisive. Regarding alignment, because the reference set covers only early-stage interactions and the alignment methods share a fixed early endpoint, this benchmark could not effectively distinguish alignment methods **(Supplementary Fig. 14)**. It therefore evaluates early-stage signaling, while the *eya* anchor assesses alignment at the spermatogonia-to-spermatocyte transition.

We further selected literature-supported pairs assigned early by DRAFT across all five seeds but late by the expression product **(Fig. 6d, Supplementary Table 9)**. These included hub-derived Upd1/2/3–Dome^41,86^, as well as cyst-cell-derived Wnt4–Arr, Wnt4– Fz/Fz2/Fz4^23^, and Gbb–Sax^42,43^. Their DRAFT scores peaked between GSC and Sp-16, consistent with the known roles of these pathways in early germ cells, whereas expression-product scores peaked at 1° Sc_3–2° Sc_5 (**Fig. 6d)**. Previously unreported pairs assigned earlier by DRAFT than by the expression product in at least four of five seeds included hub-derived Pyr–Htl, Sema1a/2a/5c–PlexA and Wnt2/Wnt5–Fz3, and cyst-cell-derived CadN–CadN. These pairs represent candidates for further experimental validation.

## Discussion

We propose DRAFT, a framework for inferring dynamic gene regulatory networks from single-cell multi-omics trajectories. DRAFT explicitly models nonlinear interactions among regulators and temporal dependencies within a unified deep graph model. Simulation benchmarks show gains in accuracy and temporal smoothness, while independent reference data support the networks inferred in mouse reprogramming and *Drosophila* adult spermatogenesis. Beyond recovering known regulatory programs, DRAFT identifies differences in the timing of high network centrality and RNA expression. For *Mondo*, germline-specific knockdown altered GSC number and expression of the candidate targets, *kuz* and *mamo*, supporting a candidate early germline role. This illustrates how DRAFT can suggest regulatory hypotheses that are not apparent from expression alone. The inferred networks further support alignment of co-developing lineages and analyses of stage-specific cell–cell communication.

Despite the improvements, the inferred relationships by DRAFT primarily capture co-variation and coordinated patterns rather than causal regulatory evidence. The inferred edges are also unsigned and do not distinguish activating from repressive relationships. Therefore, the results alone cannot determine regulatory mechanisms or directionality; key predictions still require experimental perturbation and orthogonal validation.

DRAFT also has several important limitations. First, DRAFT relies on a prior graph constructed from motifs and genomic distance, whose quality largely determines the upper bound of candidate regulatory edges. In species lacking high-quality motif data or genome annotations, this may limit the performance. Second, DRAFT only accepts integer time labels along a linear or tree-structured lineage, which requires multi-omics trajectories with clearly defined time batches or discrete stages. This may be insufficient for processes with complex branching or continuous time labels. Third, DRAFT only uses transcriptomic and chromatin-accessibility data, and other modalities, like epigenetic modifications and spatial positions, have not yet been incorporated. For cross-lineage alignment, DRAFT assumes that the lineages share the same developmental start and end times and remain in contact at corresponding stages. These strong assumptions may limit the applicability. Communication scores are calculated from ligand expression and receptor network centrality, without explicitly modelling ligand–receptor binding affinities or spatial constraints. Finally, further genetic and stage-resolved analyses will be required to validate *Mondo*’s function in the germline.

Future work can extend this study in several directions. At the model level, we may explore incorporating time-series CRISPR screens as priors, and jointly modeling them with proteomic or spatial transcriptomic data to improve causality. Meanwhile, we may use differential equation-based neural networks, thereby allowing continuous time label inputs. At the application level, DRAFT can be further applied to more complex and fine-grained dynamic processes such as tumor evolution and immune responses.

## Methods

### DRAFT model

DRAFT learns stage-resolved representations of transcription factors (TFs), regulatory elements (REs) and target genes (TGs) from single-cell RNA and chromatin-accessibility profiles ordered along a developmental trajectory, and scores candidate regulatory relationships by the similarity of these representations. A detailed mathematical description is given in the **Supplementary Information.**

### Prior graph and node features

For paired multiome data, DRAFT uses a static directed prior graph with TF, RE and TG nodes. TF→RE edges connect a TF to REs with high motif-enrichment scores, and RE→TG edges connect an RE to genes whose transcription start site (TSS) lies within a fixed distance. Each edge is complemented by a reverse edge that is used only for message passing. The same prior is used at every stage. Node features are stage-specific: TF and TG nodes carry their RNA expression and RE nodes their accessibility in the cells of that stage, zero-padded to the largest stage size so that all stages share one input dimension; padding does not pair cells across stages. For unpaired RNA and ATAC data, DRAFT uses a reduced graph of TF and TG nodes in which accessible motif-containing regions provide the evidence for candidate TF–TG edges, and only expression is used as node features.

### Architecture

At each stage, two relation-specific graph attention layers^16^ with two heads propagate information along the four edge types; each layer is followed by normalization and dropout, and a linear residual projection of the input features is added to the second layer’s output. A gated recurrent unit (GRU) then links the stages. Its hidden state starts at zero at the root stage and is passed from each stage to its child stages along the lineage tree, so that branches share all parameters but are unrolled separately. A linear skip connection of the current stage representation is added to the GRU output, and a linear layer gives the final embedding of every node at every stage.

### Training

DRAFT is trained to reconstruct the prior graph. At each stage, prior TF–RE and RE–TG edges are positives and an equal number of uniformly sampled non-edges are negatives; the edge logit is the inner product of the two node embeddings, and binary cross-entropy losses are summed over relations and stages. An optional smoothness term used for all real-data experiments penalizes the squared Frobenius distance between edge-weight matrices of each stage and its parent stage. Its weight is set at the first epoch to the ratio of the reconstruction and smoothness losses and then held fixed **(Supplementary Table 10 and 11).** Experimental reference networks and simulated ground truth were never used for training or model selection.

### Network construction

After training, each TF–TG pair is scored at each stage by the cosine similarity of the two embeddings. For *Drosophila*, candidates are restricted to genes detected in at least 5% of the cells of the stage and to the filtered TF list. Pairs above a cosine threshold θ are ranked; the K highest-scoring pairs are kept; this retains stage-specific regulators with few strong edges. In the paired model, each selected TG is linked to its two best-supported accessible REs, ranked by a weighted combination of RE–TG embedding similarity, distance to the TSS, TF–RE similarity and RE accessibility, and a TF–RE link is added when a TF and an RE share a selected target. *Drosophila* stage networks for edge clustering used θ = 0.83, K = 10,000; mouse networks used genes detected in at least 30% of cells, θ = 0.85, K = 50,000.

### Continuous networks

To obtain networks between sampled stages, the node embeddings were resampled on a grid with *k* equally spaced points between consecutive stages using a Gaussian kernel over stages (σ = 0.7 stage units) normalized to sum to one. Networks on this grid (dense networks) used *k=4*.

### Simulated data

Paired RNA and ATAC profiles were generated using scMultiSim^22^ 1.2.0 with 55 cell identity factors, diff.cif.fraction = 0.60, cif.sigma = 0.18 and native kinetic RNA simulation (do.velocity = TRUE). For the linear simulation, an explicit sampling grid provided 200 cells along each of five consecutive trajectory segments, yielding 1,000 cells per dataset. The main regulatory network comprised 50 TFs and 150 TGs assigned to three modules. Each TG had 12 candidate TFs: nine within its module and three outside. Six within-module candidates were active per TG at each stage, yielding 900 active edges among 1,800 candidate pairs. Between consecutive stages, 90 active edges (10%) were replaced by inactive within-module candidates of the same TG, preserving 6 active regulators per TG. Replacement was one-way: removed edges were not reintroduced, and newly activated edges were retained. Edge weights were binary and constant within each segment. These predefined TF–TG networks replaced scMultiSim’s random GRN update schedule. For each TG, its candidate TFs were randomly partitioned into six pairs, each assigned to a synthetic RE linked to that TG, producing 900 REs. The resulting TF– RE and RE–TG maps remained fixed across time. Latent RE accessibility was scaled by *0.2 + 0.8*, where was the fraction of assigned TFs actively regulating the linked TG. Accessibility influenced transcription through scMultiSim’s ATAC effect (atac.effect = 0.8; atac.p_zero = 0.3). Independent Poisson observation sampling at depth 1 was applied to both assays. Inactive candidates supplied false-prior relationships; no additional random edges were added. For branching simulations, a shared trunk (T0–T1) split into two daughter branches (A2–A4 and B2–B4). Networks evolved independently along each parent–child transition using the main simulation’s replacement rule, while candidate priors and RE maps remained fixed. A single scMultiSim run generated 200 cells per stage; other settings matched the linear simulation.

### Mouse reprogramming data

Following the scMTNI pipeline^13^, we analysed unpaired scATAC-seq (GSE208620^13^) and scRNA-seq (GSE108222^24^) profiles of mouse embryonic fibroblast (MEF) reprogramming at day 0, days 3, 6, 9 and 12 and the embryonic stem cell (mESC) endpoint. ATAC fragments were counted in non-overlapping 1-kb bins of mm10 and bins were assigned to the nearest gene by TSS distance (30,344 nuclei after keeping the published high-quality barcodes). RNA counts were normalized to 10,000 counts per cell and log1p-transformed; genes detected in fewer than five cells at a sampling time were removed. The 11,926 genes shared by both modalities were integrated with LIGER^87^ (rliger 0.5.1, *k* = 8), and a minimum spanning tree over cluster pseudobulk profiles defined the lineage, with a successful (C4–C8–C1–C5) and an unsuccessful (C4–C8–C7–C3–C2) branch. Cluster-specific TF–TG priors were generated with the scMTNI prior-generation pipeline^13^: CisBP motif instances^88^ in each cluster’s ATAC peaks linked a TF to genes whose TSS lay within 5 kb of the peak, and edges were scored by percentile rank. Rerunning the pipeline reproduced the priors released with that study, and DRAFT used edges with a score above 0.1.

### *Drosophila* testis multiome data

Paired snRNA-seq and snATAC-seq profiles of the adult *Drosophila melanogaster* testis apex (GSE277132^23^) were reprocessed (10,335 nuclei). RNA was processed with scanpy^89^ 1.10.1 (filtering on detected genes, counts and mitochondrial fraction; Scrublet^90^ doublet removal; library-size normalization and log transformation; Harmony^91^ integration across samples). ATAC peaks were called with MACS3^92^, restricted to nuclei passing RNA quality control, filtered on fragment counts, fraction of reads in peaks, TSS enrichment and nucleosome signal, and embedded by TF–IDF, latent semantic indexing and Harmony. Annotations from the source study defined 15 germline stages (GSC, Gb, Sp-2, Sp-4, Sp-8, Sp-16, 1° Sc_1–3 and 2° Sc_1–6) and nine somatic groups (CySC, eSCC-1–3, lSCC-1–3 and the two terminal branches SdA and SdB); hub cells were analysed separately. Peaks were used as REs. TF–RE prior edges were the highest 3% ranked motif–region pairs from SCENIC+/cisTarget^1^ with a *Drosophila* motif collection, and RE–TG edges linked peaks to TSSs within 100 kb in dm6. The prior had 761 TF nodes with motif annotations; network reconstruction was restricted to a filtered list of 272 TFs by FlyBase^93^. Separate models were trained for the germline and somatic lineages.

### Baseline methods

Baselines were run from pinned upstream versions using their native training procedures. Method-specific settings were held fixed across the reported benchmark runs. Reference-network and simulated ground-truth labels were not used for model fitting. Where a method’s own prior construction could not be applied, the DRAFT prior was supplied instead. For the simulation the constructed prior graph was supplied to all methods.

#### Simulation

We used SCENIC+’s TF2G module (1.0a2) to infer gradient-boosting importances from log-normalized RNA. The full eRegulon workflow was not applied because the simulated regulatory regions lack the genomic sequences required for motif analysis. scMTNI (commit c4d4850) treated the stages as a lineage chain and used the prior-connected TF–TG pairs as a soft prior (β0 = −3, β1 = 2). Its edge scores are selection frequencies over 50 half-cell subsamples. Dictys (1.1.0) fitted its Ornstein– Uhlenbeck model to raw RNA counts at each stage (4,000 steps), with the prior-connected pairs as the allowed-edge mask. scDoRI (1.0.0) was trained on paired RNA and ATAC with the TF–RE and RE–TG priors (40 topics, native early stopping). Random scores were drawn from Uniform(0, 1).

#### Mouse

SCENIC (pySCENIC 0.12.1) was run per cluster, with GRNBoost2 and cisTarget pruning against the mm10 mc9nr rankings. Dictys (1.1.0) was also run per cluster on the original UMI counts, with the DRAFT prior as the edge mask. In clusters with fewer cells than regulators, its covariance rank was capped at the number of cells minus one. For scMTNI, the published scMTNI+Prior networks were used. scDoRI required paired data and was not run.

#### Drosophila

scMTNI and Dictys inferred TF–TG networks only, so the RE–TG benchmark compared DRAFT with SCENIC+ and scDoRI. Both used the same cells, peaks and RE– TG prior as DRAFT. The SCENIC+ region-to-gene module was refitted on both lineages: topics and imputed accessibility (pycisTopic 2.0a, 40 topics) were computed once across all cells, and the gradient-boosting regression was then fitted separately on the cells of each stage. scDoRI (1.0.0) used the DRAFT TF–RE prior in place of motif scanning, with 40 topics. Its stage RE–TG score is the stage-mean accessibility of the peak multiplied by the learned gene–peak weight. At each stage, the random control sampled as many TF–RE and RE–TG edges as the prior network contains.

### Benchmark evaluation

#### Simulation

At each stage, all pairs in the fixed TF–TG candidate set were scored, with active edges as positives and inactive candidates as negatives, and the area under the receiver operating characteristic curve (AUROC) was averaged over stages without thresholding or negative subsampling. Replication numbers and the units used to calculate error bars are specified in the corresponding figure legends.

#### Mouse

Networks inferred for each of seven cell clusters were evaluated against the three mESC gold-standard networks released with the original scMTNI study^13^: ChIP-based interactions from ESCAPE and ENCODE, perturbation-based interactions from ESCAPE and Nishiyama et al., and their intersection. We followed the original TF/target filtering and candidate-pair definition, evaluating gold-covered TF–TG combinations within each cluster’s available genes and regulators, excluding self-pairs. Missing prediction scores were assigned zero. AUPR was calculated using the official preparation script and AUCCalculator implementation at natural class prevalence. To avoid the fixed Top-1,000-edge cutoff of the original F1 benchmark, we instead calculated AUROC using uniform 1:1 positive-to-negative sampling, evaluated on a fixed half-per-class test subset. Results summarize the mean and sample SD across seven cell clusters.

#### Drosophila

RE–TG predictions were compared with stage-matched testis Hi-C loops (GSE156567^45^) at 2-kb resolution: a TG–peak pair is positive when one loop anchor overlaps the TG TSS and the other the peak. Spermatogonial loops were evaluated at Sp-2 to Sp-16 and spermatocyte loops at 1° Sc_1 to 2° Sc_6. Evaluation was restricted to the shared RE–TG prior, with an equal number of randomly drawn prior pairs without Hi-C support as negatives; AUROC was the primary metric and each stage contributed one value.

#### Temporal smoothness

For each method, the stage-specific TF–TG edge-weight matrices were restricted to the same candidate pairs, pairwise Frobenius distances were computed, and smoothness was defined as one minus the mean distance between adjacent stages divided by the largest pairwise distance. Smoothness describes how gradually predicted networks change; it does not measure accuracy or absolute changes and a time-invariant prediction attains the maximum.

### Edge clustering and topic modelling

For edge clustering, the stage networks were combined into a binary edge-by-stage matrix (1 where an edge was selected at a stage, 0 otherwise), standardized per stage and clustered with k-means into 12 edge clusters. Within each edge cluster, TFs were ranked by the number of distinct targets (centrality; this is network degree, not TF abundance or binding). For topic modelling, each TF–TG edge was treated as a document whose words are the stages at which the edge is among the top-scoring edges, and latent Dirichlet allocation was fitted with 12 topics for Drosophila and 10 for mouse; a topic’s activity across stages is its topic–stage weight, normalized to sum to one over stages. For *Drosophila*, documents were built from the 10,000 highest-scoring TF–TG pairs of each stage (TFs, REs and TGs detected in at least 5% of the stage’s cells), and the genes targeted by the 500 highest-weighted edges of each topic were tested with FlyEnrichr^94^ (GO_Biological_Process_2018). Mouse topics were tested with Enrichr^95^ (GO Biological Process 2021) against the 7,764 network targets. Benjamini–Hochberg adjusted *P* values are reported. Edge-cluster and topic numbers in figures and tables refer to a single frozen run.

### Cross-lineage alignment

At each grid point, the weighted TF–TG adjacency of the dense network (genes present in both lineages) was degree-normalized and summarized by a fingerprint of its 20 leading singular values (z-scored across time). This alignment seeks stages with similar global regulatory-network organizations across lineages. Restricting the comparison to shared TFs and TGs reduces differences in gene coverage. Dynamic time warping^77^ matched somatic to germline grid points using the squared Euclidean distance between their standardized spectral fingerprints as the alignment cost, within a band of ±6 grid points. The resulting warp was smoothed with a curvature penalty, made monotone by isotonic regression, computed once and stored. SdA and SdB were treated as one terminal stage placed at the last germline stage. A linear alignment served as the baseline. As an independent check, the largest rise of *eya* in the soma and of *kmg*, *aly* and *can* in the germline was located in each lineage (support from 200 bootstrap resamples of cells) and the somatic switch was mapped onto germline stages by each alignment.

### Ligand–receptor timing

Candidates were FlyPhoneDB^83^ ligand–receptor pairs with both genes in the networks (171 pairs), scored for hub→germline, hub→soma, soma→germline and germline→soma. A pair was kept when the ligand was expressed in ≥5% of the cells of a sender group and the receptor in ≥5% of the cells of a receiver stage (81, 83, 74 and 84 pairs). The ligand term is the mean ligand expression in the sender (constant for the hub; for somatic and germline senders, read at the aligned position of each receiver stage). The receptor network centrality is the receptor’s percentile among the cosine similarities of all active TF–TG pairs at the stage, averaged over active TFs. Their product is the score of the pair at a receiver stage, set to zero where the receptor is detected in fewer than 5% of receiver cells. Expression-based comparators — the ligand × receptor expression product, CellChat^84^ and LIANA+^85^ — used the same pairs, alignment and receptor gate. A pair was called early if its scores peaked at GSC–Sp-16 (germline) or CySC–eSCC-3 (soma), late otherwise. Its earliness score was (early maximum − late maximum) / overall maximum. Literature pairs with a stated sender and receiver in adult testis signalling studies **(Supplementary Table 8)** were compared with all other candidates of the same direction by the AUROC of the earliness score, averaged over 1,000 size-matched draws of non-literature pairs.

CellChat and LIANA+ were run on the same dataset as DRAFT, using log-normalized RNA and 25 groups (the hub, nine somatic and fifteen germline stages). Both used the same fly ligand–receptor list of 171 single-gene pairs, without complex or cofactor annotations. For CellChat, the inference functions of GitHub commit 75253cd (version 2.2.0.9001) were applied unchanged. The computeCommunProb function used a 10% truncated mean, Kh = 0.5, n = 1 and no population-size weighting. The communication probability of every sender–receiver group pair was used, without the permutation significance filter. LIANA+ (1.10.1.dev0, commit 08590db) was run with rank_aggregate, which combines CellPhoneDB, Connectome, log2FC, NATMI and SingleCellSignalR by robust rank aggregation (at least 5 cells per group). Its expression-proportion filter was turned off (expr_prop = 0), so the shared 5% receptor gate was the only expression filter.

### Fly husbandry and germline knockdown

Flies were raised on standard cornmeal food containing barley malt, molasses, yeast extract, soy flour, propionic acid and Nipagin at 25 °C. Strains are listed in **Supplementary Table 12**. For germline-specific knockdown, *nos-Gal4::VP16* virgin females were crossed with *UAS-Mondo* RNAi males; controls were *nos-Gal4::VP16* females crossed with *UAS-mCherry* RNAi males. Crosses were raised at 18 °C to reduce GAL4 activity during development and progeny were shifted to 29 °C at eclosion for 7 days before analysis, alongside age-matched controls.

### smFISH and immunostaining

smFISH combined with immunostaining was performed as described in the original study^23^, with minor modifications. Testes were dissected in ice-cold 1× PBS, fixed in 4% paraformaldehyde for 20 min at room temperature and washed three times for 10 min in 0.2% PBT (1× PBS, 0.2% Triton X-100, 0.2% Tween-20). Samples were blocked for 30 min at 37 °C and 500 r.p.m. in 2× SSC, 10% dextran sulfate, 1 mg ml−1 tRNA, 2 mM vanadyl ribonucleoside complex (NEB, S142) and 0.5% BSA, then incubated overnight at 37 °C and 800 r.p.m. with smFISH probes (1:100) and primary antibodies. After three 10-min washes in 0.2% PBT, samples were incubated with secondary antibodies in 0.2% PBT for 2 h, washed three times and mounted in Vectashield (Vector Laboratories, H1000). Antibodies and probes are listed in **Supplementary Table 13**. Images were acquired on a Leica TCS SP8 confocal microscope.

### Image quantification

GSCs were counted with a two-step three-dimensional segmentation: the Fas3-labelled hub was segmented with a custom 3D U-Net and the germline with a custom Cellpose^96^ Vasa model, and Vasa-positive cells contacting the hub were counted as GSCs. *Mondo*, *mamo* and *kuz* smFISH intensities were measured as the mean intensity within each segmented GSC. Models and analysis code are available on request.

### Statistics

The number of hub-contacting germ cells per testis was compared between genotypes with a two-sided Mann–Whitney U test (n = testes). For smFISH quantification, fluorescence intensities were measured in individual germline stem cells (GSCs), and distributions were compared between genotypes using a two-sided Mann–Whitney U test on individual GSC measurements. Numbers of GSCs, testes and independent experiments are given in the figure legends. Analyses used Python. P < 0.05 was considered significant; exact P values and effect sizes are reported in the figure legends and Source Data.

## Data Availability

For mouse reprogramming, the scATAC-seq data were downloaded from the Gene Expression Omnibus (GEO) under accession **GSE208620**, and the scRNA-seq data for the same time points were downloaded from GEO under accession **GSE108222**. The mESC ChIP-seq data were downloaded from the scMTNI source data (https://zenodo.org/records/7879228). The perturbation data were downloaded from the Escape database (http://www.maayanlab.net/ESCAPE). For *Drosophila* testis development data, we collected paired multiome data from GEO under accession **<u>GSE277132</u>**. The Hi-C data on SpG and SpC phases were downloaded from GEO under accession **GSE156567**. The collection of existing ligand-receptor datasets between the two lineages used for predicting cell-cell communication is derived from the original study (https://dawa.cos.uni-heidelberg.de). All data used in this paper as input and model checkpoints can be downloaded from Zenodo (10.5281/zenodo.23678662).

## Code Availability

DRAFT is available from GitHub (https://github.com/velten-group/DRAFT.git). Code to reproduce the results in the manuscript can be found on GitHub (https://github.com/velten-group/DRAFT-analysis.git).

## Contributions

Zehua Zhang designed the computational model and performed all analyses. Pallavi Santhi Sekhar designed and carried out the experimental work in *Drosophila*. Patrick van Nierop y Sanchez, as an author of the original *Drosophila* dataset, provided data support and valuable scientific input. Ingrid Lohmann supervised the biological interpretation and experimental validation of the *Drosophila* study. Britta Velten supervised the overall project.

## Acknowledgements

We thank Prof. Dr. Carl Herrmann and Wangjun Hu for critical and constructive suggestions, and we are grateful to Nikolai Köhler, Thomas Greulich and Jana Braunger for their technical assistance. We thank Yuqing Hong for constructive suggestions on the biological interpretation. We acknowledge the data storage service SDS@hd supported by the Ministry of Science, Research and the Arts Baden-Württemberg (MWK) and the German Research Foundation (DFG) through grant INST 35/1503-1 FUGG. The work was funded by the DFG (project 530144743).

## References

1. Bravo González-Blas, C., et al. SCENIC+: single-cell multiomic inference of enhancers and gene regulatory networks. Nat. Methods 20, 1355–1367 (2023).

2. Tanay, A. & Regev, A. Scaling single-cell genomics from phenomenology to mechanism. Nature 541, 331–338 (2017).

3. Smith, Z. D., Sindhu, C. & Meissner, A. Molecular features of cellular reprogramming and development. Nat. Rev. Mol. Cell Biol. 17, 139–154 (2016).

4. Aibar, S. et al. SCENIC: single-cell regulatory network inference and clustering. Nat. Methods 14, 1083–1086 (2017).

5. Badia-i-Mompel, P. et al. Gene regulatory network inference in the era of single-cell multi-omics. Nat. Rev. Genet. 24, 739–754 (2023).

6. Cao, Z.-J. & Gao, G. Multi-omics single-cell data integration and regulatory inference with graph-linked embedding. Nat. Biotechnol. 40, 1458–1466 (2022).

7. Fleck, J. S. et al. Inferring and perturbing cell fate regulomes in human brain organoids. Nature 621, 365–372 (2023).

8. Kartha, V. K. et al. Functional inference of gene regulation using single-cell multi-omics. Cell Genomics 2, 100166 (2022).

9. Duren, Z., Chen, X., Jiang, R., Wang, Y. & Wong, W. H. Modeling gene regulation from paired expression and chromatin accessibility data. Proc. Natl. Acad. Sci. 114, E4914–E4923 (2017).

10. Kamimoto, K. et al. Dissecting cell identity via network inference and in silico gene perturbation. Nature 614, 742–751 (2023).

11. Wang, L. et al. Dictys: dynamic gene regulatory network dissects developmental continuum with single-cell multiomics. Nat. Methods 20, 1368–1378 (2023).

12. Saraswat, M. et al. Decoding Plasticity Regulators and Transition Trajectories in Glioblastoma with Single-cell Multiomics. 2025.05.13.653733 Preprint at 10.1101/2025.05.13.653733 (2025).

13. Zhang, S. et al. Inference of cell type-specific gene regulatory networks on cell lineages from single cell omic datasets. Nat. Commun. 14, 3064 (2023).

14. Lin, Y. et al. Data integration and inference of gene regulation using single-cell temporal multimodal data with scTIE. Genome Res. 34, 119–133 (2024).

15. Kipf, T. N. & Welling, M. Semi-Supervised Classification with Graph Convolutional Networks. Preprint at 10.48550/arXiv.1609.02907 (2017).

16. Veličković, P., et al. Graph Attention Networks. Preprint at 10.48550/arXiv.1710.10903 (2018).

17. Hamilton, W. L., Ying, R. & Leskovec, J. Inductive Representation Learning on Large Graphs. Preprint at 10.48550/arXiv.1706.02216 (2018).

18. Ma, A. et al. Single-cell biological network inference using a heterogeneous graph transformer. Nat. Commun. 14, 964 (2023).

19. Hochreiter, S. & Schmidhuber, J. Long Short-Term Memory. Neural Comput. 9, 1735–1780 (1997).

20. Cho, K. et al. Learning Phrase Representations using RNN Encoder–Decoder for Statistical Machine Translation. in Proceedings of the 2014 Conference on Empirical Methods in Natural Language Processing (EMNLP) (eds Moschitti, A., Pang, B. & Daelemans, W.) 1724–1734 (Association for Computational Linguistics, Doha, Qatar, 2014). doi:10.3115/v1/D14-1179.

21. Hasanaj, E., Póczos, B. & Bar-Joseph, Z. Recovering time-varying networks from single-cell data. Bioinformatics 41, i628–i636 (2025).

22. Li, H., Zhang, Z., Squires, M., Chen, X. & Zhang, X. scMultiSim: simulation of single-cell multi-omics and spatial data guided by gene regulatory networks and cell-cell interactions. Nat. Methods 22, 982–993 (2025).

23. van Nierop y Sanchez, P., et al. Dissecting the enhancer gene regulatory network in early Drosophila spermatogenesis. Nat. Commun. 16, 6766 (2025).

24. Tran, K. A. et al. Defining Reprogramming Checkpoints from Single-Cell Analyses of Induced Pluripotency. Cell Rep. 27, 1726–1741.e5 (2019).

25. The ENCODE Project Consortium. An Integrated Encyclopedia of DNA Elements in the Human Genome. Nature 489, 57–74 (2012).

26. Xu, H. et al. ESCAPE: database for integrating high-content published data collected from human and mouse embryonic stem cells. Database J. Biol. Databases Curation 2013, bat045 (2013).

27. Nishiyama, A. et al. Uncovering Early Response of Gene Regulatory Networks in ESCs by Systematic Induction of Transcription Factors. Cell Stem Cell 5, 420–433 (2009).

28. Chronis, C. et al. Cooperative binding of transcription factors orchestrates reprogramming. Cell 168, 442–459.e20 (2017).

29. Tavares, A. H. et al. TGF-β1-dependent expression of FOXS1 attenuates adipogenic potential and enhances a myofibroblast cellular phenotype. J. Biol. Chem. 301, 110563 (2025).

30. Nefzger, C. M. et al. Cell Type of Origin Dictates the Route to Pluripotency. Cell Rep. 21, 2649–2660 (2017).

31. Jeon, H. et al. Comprehensive Identification of Krüppel-Like Factor Family Members Contributing to the Self-Renewal of Mouse Embryonic Stem Cells and Cellular Reprogramming. PLoS ONE 11, e0150715 (2016).

32. Festuccia, N. et al. Esrrb Is a Direct Nanog Target Gene that Can Substitute for Nanog Function in Pluripotent Cells. Cell Stem Cell 11, 477–490 (2012).

33. Yu, H., Kunarso, G., Hong, F. H. & Stanton, L. W. Zfp206, Oct4, and Sox2 Are Integrated Components of a Transcriptional Regulatory Network in Embryonic Stem Cells. J. Biol. Chem. 284, 31327–31335 (2009).

34. Kwon, Y.-W. et al. E-Ras improves the efficiency of reprogramming by facilitating cell cycle progression through JNK–Sp1 pathway. Stem Cell Res. 15, 481–494 (2015).

35. Baksh, S. S., Pratt, R. E., Gomez, J., Dzau, V. J. & Hodgkinson, C. P. A novel Cbx1, PurB, and Sp3 complex mediates long-term silencing of tissue- and lineage-specific genes. J. Biol. Chem. 298, 102053 (2022).

36. Zhou, C., Wang, M., Zhang, C. & Zhang, Y. The transcription factor GABPA is a master regulator of naive pluripotency. Nat. Cell Biol. 27, 48–58 (2025).

37. Marión, R. M. et al. A p53-mediated DNA damage response limits reprogramming to ensure iPS cell genomic integrity. Nature 460, 1149–1153 (2009).

38. Marion, R. M. et al. Telomeres Acquire Embryonic Stem Cell Characteristics in Induced Pluripotent Stem Cells. Cell Stem Cell 4, 141–154 (2009).

39. Li, R. et al. A mesenchymal-to-epithelial transition initiates and is required for the nuclear reprogramming of mouse fibroblasts. Cell Stem Cell 7, 51–63 (2010).

40. Li, Z., Dang, J., Chang, K.-Y. & Rana, T. M. MicroRNA-mediated regulation of extracellular matrix formation modulates somatic cell reprogramming. RNA 20, 1900–1915 (2014).

41. Tulina, N. & Matunis, E. Control of Stem Cell Self-Renewal in Drosophila Spermatogenesis by JAK-STAT Signaling. Science 294, 2546–2549 (2001).

42. Kawase, E., Wong, M. D., Ding, B. C. & Xie, T. Gbb/Bmp signaling is essential for maintaining germline stem cells and for repressing bam transcription in the Drosophila testis. Development 131, 1365–1375 (2004).

43. Shivdasani, A. A. & Ingham, P. W. Regulation of stem cell maintenance and transit amplifying cell proliferation by tgf-beta signaling in Drosophila spermatogenesis. Curr. Biol. CB 13, 2065–2072 (2003).

44. Zoller, R. & Schulz, C. The Drosophila cyst stem cell lineage: Partners behind the scenes? Spermatogenesis 2, 145–157 (2012).

45. Ilyin, A. A. et al. Comparison of genome architecture at two stages of male germline cell differentiation in Drosophila. Nucleic Acids Res. 50, 3203–3225 (2022).

46. Angulo, B. et al. DREF Genetically Counteracts Mi-2 and Caf1 to Regulate Adult Stem Cell Maintenance. PLoS Genet. 15, e1008187 (2019).

47. Hayashi, M. et al. Conserved role of Ovo in germline development in mouse and Drosophila. Sci. Rep. 7, 40056 (2017).

48. Streit, A., Bernasconi, L., Sergeev, P., Cruz, A. & Steinmann-Zwicky, M. mgm 1, the earliest sex-specific germline marker in Drosophila, reflects expression of the gene esg in male stem cells. Int. J. Dev. Biol. 46, 159–166 (2002).

49. Voog, J. et al. Escargot restricts niche cell to stem cell conversion in the Drosophila testis. Cell Rep. 7, 722–734 (2014).

50. Kong, R. et al. A regulatory loop of JAK/STAT signalling and its downstream targets represses cell fate conversion and maintains male germline stem cell niche homeostasis. Cell Prolif. 57, e13648 (2024).

51. Havula, E. et al. Mondo/ChREBP-Mlx-Regulated Transcriptional Network Is Essential for Dietary Sugar Tolerance in Drosophila. PLOS Genet. 9, e1003438 (2013).

52. Xing, Y., Su, T. T. & Ruohola-Baker, H. Tie-mediated signal from apoptotic cells protects stem cells in Drosophila melanogaster. Nat. Commun. 6, 7058 (2015).

53. Chen, W. et al. CG8005 Mediates Transit-Amplifying Spermatogonial Divisions via Oxidative Stress in Drosophila Testes. Oxid. Med. Cell. Longev. 2020, 2846727 (2020).

54. Khan, C., Rusan, N. M. & Baker, N. E. Dual contributions of Xrp1 to genome integrity through the DNA damage response and cell competition. PLoS Genet. 22, e1012309 (2026).

55. Ochi, N. et al. Cell competition is driven by Xrp1-mediated phosphorylation of eukaryotic initiation factor 2α. PLOS Genet. 17, e1009958 (2021).

56. Li, Y., Ma, Q., Cherry, C. M. & Matunis, E. L. Steroid signaling promotes stem cell maintenance in the Drosophila testis. Dev. Biol. 394, 129–141 (2014).

57. Issigonis, M. & Matunis, E. The Drosophila BCL6 homolog ken and barbie promotes somatic stem cell self-renewal in the testis niche. Dev. Biol. 368, 181–192 (2012).

58. Ma, Q., Wawersik, M. & Matunis, E. L. The Jak-STAT target Chinmo prevents sex transformation of adult stem cells in the Drosophila testis niche. Dev. Cell 31, 474– 486 (2014).

59. Ables, E. T., Bois, K. E., Garcia, C. A. & Drummond-Barbosa, D. Ecdysone response gene *E78* controls ovarian germline stem cell niche formation and follicle survival in *Drosophila*. Dev. Biol. 400, 33–42 (2015).

60. Butsch, T. J., Johnson, A. E. & Bohnert, K. A. Meiotic activation of Mitf/TFEB declines with age in the Drosophila testis. MicroPublication Biol. 10.17912/micropub.biology.001509 (2025).

61. Sanchez, C. G. et al. Regulation of Ribosome Biogenesis and Protein Synthesis Controls Germline Stem Cell Differentiation. Cell Stem Cell 18, 276–290 (2016).

62. Sênos Demarco, R., Uyemura, B. S., D’Alterio, C. & Jones, D. L. Mitochondrial fusion regulates lipid homeostasis and stem cell maintenance in the Drosophila testis. Nat. Cell Biol. 21, 710–720 (2019).

63. White-Cooper, H. Molecular mechanisms of gene regulation during Drosophila spermatogenesis. Reproduction 139, 11–21 (2010).

64. Benner, L., Muron, S., Gomez, J. G. & Oliver, B. OVO Positively Regulates Essential Maternal Pathways by Binding Near the Transcriptional Start Sites in the Drosophila Female Germline. bioRxiv 2023.11.01.565184 (2024) doi:10.1101/2023.11.01.565184.

65. Mattila, J. et al. Mondo-Mlx Mediates Organismal Sugar Sensing through the Gli-Similar Transcription Factor Sugarbabe. Cell Rep. 13, 350–364 (2015).

66. McLeod, C. J., Wang, L., Wong, C. & Jones, D. L. Stem Cell Dynamics in Response to Nutrient Availability. Curr. Biol. 20, 2100–2105 (2010).

67. Yang, H. & Yamashita, Y. M. The regulated elimination of transit-amplifying cells preserves tissue homeostasis during protein starvation in Drosophila testis. Development 142, 1756–1766 (2015).

68. Huang, W.-H. et al. Excess dietary sugar impairs Drosophila adult stem cells via elevated reactive oxygen species-induced JNK signaling. Development 151, dev201772 (2024).

69. Clémot, M., D’Alterio, C., Kwang, A. C. & Jones, D. L. mTORC1 is required for differentiation of germline stem cells in the Drosophila melanogaster testis. PLOS ONE 19, e0300337 (2024).

70. Pan, D. & Rubin, G. M. Kuzbanian Controls Proteolytic Processing of Notch and Mediates Lateral Inhibition during Drosophila and Vertebrate Neurogenesis. Cell 90, 271–280 (1997).

71. Ng, C. L., Qian, Y. & Schulz, C. Notch and Delta are required for survival of the germline stem cell lineage in testes of Drosophila melanogaster. PLOS ONE 14, e0222471 (2019).

72. Nakamura, S. et al. A truncated form of a transcription factor Mamo activates vasa in Drosophila embryos. *Commun*. Biol. 2, 422 (2019).

73. de Cuevas, M. & Matunis, E. L. The stem cell niche: lessons from the Drosophila testis. Dev. Camb. Engl. 138, 2861–2869 (2011).

74. Fairchild, M. J., Smendziuk, C. M. & Tanentzapf, G. A somatic permeability barrier around the germline is essential for Drosophila spermatogenesis. Development 142, 268–281 (2015).

75. Alpert, A., Moore, L. S., Dubovik, T. & Shen-Orr, S. S. Alignment of single-cell trajectories to compare cellular expression dynamics. Nat. Methods 15, 267–270 (2018).

76. Sugihara, R., Kato, Y., Mori, T. & Kawahara, Y. Alignment of single-cell trajectory trees with CAPITAL. Nat. Commun. 13, 5972 (2022).

77. Sakoe, H. & Chiba, S. Dynamic programming algorithm optimization for spoken word recognition. IEEE Trans. Acoust. Speech Signal Process. 26, 43–49 (1978).

78. Fabrizio, J. J., Boyle, M. & DiNardo, S. A somatic role for eyes absent (eya) and sine oculis (so) in Drosophila spermatocyte development. Dev. Biol. 258, 117–128 (2003).

79. Raz, A. A. et al. Emergent dynamics of adult stem cell lineages from single nucleus and single cell RNA-Seq of Drosophila testes. eLife 12, e82201 (2023).

80. Kim, J. et al. Blocking promiscuous activation at cryptic promoters directs cell type– specific gene expression. Science 356, 717–721 (2017).

81. White-Cooper, H., Leroy, D., MacQueen, A. & Fuller, M. T. Transcription of meiotic cell cycle and terminal differentiation genes depends on a conserved chromatin associated protein, whose nuclear localisation is regulated. Development 127, 5463–5473 (2000).

82. Hiller, M. A., Lin, T.-Y., Wood, C. & Fuller, M. T. Developmental regulation of transcription by a tissue-specific TAF homolog. Genes Dev. 15, 1021–1030 (2001).

83. Liu, Y. et al. FlyPhoneDB: an integrated web-based resource for cell-cell communication prediction in Drosophila. Genetics 220, iyab235 (2022).

84. Jin, S. et al. Inference and analysis of cell-cell communication using CellChat. Nat. Commun. 12, 1088 (2021).

85. Dimitrov, D. et al. LIANA+ provides an all-in-one framework for cell–cell communication inference. Nat. Cell Biol. 26, 1613–1622 (2024).

86. Kiger, A. A., Jones, D. L., Schulz, C., Rogers, M. B. & Fuller, M. T. Stem Cell Self-Renewal Specified by JAK-STAT Activation in Response to a Support Cell Cue. Science 294, 2542–2545 (2001).

87. Welch, J. D. et al. Single-Cell Multi-omic Integration Compares and Contrasts Features of Brain Cell Identity. Cell 177, 1873–1887.e17 (2019).

88. Weirauch, M. T. et al. Determination and inference of eukaryotic transcription factor sequence specificity. Cell 158, 1431–1443 (2014).

89. Wolf, F. A., Angerer, P. & Theis, F. J. SCANPY: large-scale single-cell gene expression data analysis. Genome Biol. 19, 15 (2018).

90. Wolock, S. L., Lopez, R. & Klein, A. M. Scrublet: Computational Identification of Cell Doublets in Single-Cell Transcriptomic Data. Cell Syst. 8, 281–291.e9 (2019).

91. Korsunsky, I. et al. Fast, sensitive and accurate integration of single-cell data with Harmony. Nat. Methods 16, 1289–1296 (2019).

92. Doherty, P. et al. MACS3: A Peak-calling Platform for Bulk and Single-cell Regulatory Genomics. Genomics Proteomics Bioinformatics qzag097 (2026) doi:10.1093/gpbjnl/qzag097.

93. Öztürk-Çolak, A. et al. FlyBase: updates to the Drosophila genes and genomes database. Genetics 227, iyad211 (2024).

94. Kuleshov, M. V. et al. modEnrichr: a suite of gene set enrichment analysis tools for model organisms. Nucleic Acids Res. 47, W183–W190 (2019).

95. Kuleshov, M. V. et al. Enrichr: a comprehensive gene set enrichment analysis web server 2016 update. Nucleic Acids Res. 44, W90–97 (2016).

96. Stringer, C., Wang, T., Michaelos, M. & Pachitariu, M. Cellpose: a generalist algorithm for cellular segmentation. Nat. Methods 18, 100–106 (2021).

