## Supplementary Information for "DRAFT: graph-recurrent learning of dynamic gene regulatory networks along single-cell multi-omics trajectories"

### Supplementary Methods

These Supplementary Methods give the mathematical formulation of DRAFT and of the analyses reported in the Methods. DRAFT is described once: a graph encoder on a typed prior graph, a recurrent unit along the lineage tree, and a loss that reconstructs the prior. The paired model (*Drosophila* testis and simulation) and the unpaired model (mouse reprogramming) are instances of this formulation that differ in the node types, the pair score used in the loss and the smoothness term. Data sources, cell annotation and the experimental procedures are described in the **Methods**.

#### 1 Notation and model inputs

**Lineage tree.** Each dataset is a set of developmental units $t \in T$ (stages, clusters or simulated stages, all called stages below) arranged in a rooted tree. $pa(t)$ is the parent of $t$ and $pa(t) = \varnothing$ at the root, so a linear trajectory is the special case $pa(t) = t - 1$. The germline (15 stages, GSC to *2*° Sc_6) and the simulation (five stages) are chains. The nine somatic groups form a chain from CySC to ISCC-3 that ends in two children, SdA and SdB. The seven mouse clusters form two branches from a shared root, C4–C8–C1–C5 (successful) and C4–C8–C7–C3–C2 (unsuccessful reprogramming). $C_t$ is the set of cells of stage $t$, $n_t = |C_t|$ and $n_{max} = max_t n_t$.

**Typed prior graph.** DRAFT works on a static graph with typed nodes and typed edges. The node types are TF, RE and TG in the paired model and TF and TG in the unpaired model; $V_\tau$ is the node set of type $\tau$ and $N_\tau = |V_\tau|$. The prior is a set of binary adjacency matrices $A^\varrho$, one for each prior relation $\varrho \in R_0$, with $R_0 = \{TF \to RE,\ RE \to TG\}$ in the paired model and $R_0 = \{TF \to TG\}$ in the unpaired model; we write $A^{TR}$, $A^{RG}$ and $A^{FG}$. Message passing uses the prior relations and their reverses, $R = R_0 \cup R_0^\top$; the reverse relations never enter the loss. In the paired model, the same prior is used at every stage. In both models the TF–TG candidates are the pairs joined by prior edges,

$$M = 1\left[A^{TR}A^{RG} > 0\right] \text{ (paired)}, \qquad M = A^{FG} \text{ (unpaired)}. \qquad (1)$$

In *Drosophila*, $A^{TR}$ holds the top 3% of motif–region pairs and $A^{RG}$ links each peak to the genes whose TSS lies within 100 kb; other motif-enrichment methods can supply $A^{TR}$ in the same way. In the mouse, $A^{FG}$ of cluster $t$ denotes the TF–TG input graph derived from the shared candidate prior using that cluster's ATAC accessibility cluster's prior network (edges with score above 0.1), so the mouse prior differs between clusters (Section 7). The simulated prior is constructed in Section 6.

**Node features.** A node is described at stage $t$ by its values in the cells of the stage, padded with zeros to a common length,

$$x_v^{(t)} = [(u_{vc})_{c \in C_t} \parallel 0_{n_{max} - n_t}] \in R^{n_{max}}, \qquad (2)$$

where $u_{vc}$ is the expression of a TF or TG node, or the accessibility of an RE node, in cell $c$ after a dataset-specific transform: log-normalized RNA and raw peak counts in *Drosophila*, per-gene z-scores of $log(1 + x)$ within the cluster in the mouse, and $log(1 + x)$ of counts per $10^4$ in the simulation. Within a stage, column $c$ refers to the same cell for every node: in the paired data the RNA of TF and TG nodes and the accessibility of RE nodes are measured in the same nucleus, so the features of different nodes are aligned cell by cell and carry their co-variation across the cells of the stage.

#### 2 Graph encoder[1]

The encoder maps the graph of stage $t$, with features $X^{(t)} = [x_v^{(t)}]_v$, to node representations $z_v^{(t)}$; its parameters are shared by all stages. It consists of two layers of relation-specific graph attention, each followed by normalization and dropout, and a linear residual of the input.

**Relation-specific attention.** For relation $r \in R$, head $q = 1, ..., H$, layer $l = 1,2$ and target node $v$ with in-neighbours $N_r(v)$ (no self-loops), starting from $h_v^{(0)} = x_v^{(t)}$ without an input projection,

$$\tilde{h}_u^{(l,r,q)} = \Theta^{(l,r,q)} h_u^{(l-1)}, \qquad \Theta^{(l,r,q)} \in R^{d \times d_{l-1}}, \qquad (3)$$

$$e_{uv}^{(l,r,q)} = LeakyReLU\left(a_{src}^{(l,r,q)\top} \tilde{h}_u^{(l,r,q)} + a_{dst}^{(l,r,q)\top} \Theta^{(l,r,q)} h_v^{(l-1)}\right), \qquad (4)$$

$$\alpha_{uv}^{(l,r,q)} = \frac{exp e_{uv}^{(l,r,q)}}{\sum_{u' \in N_r(v)} exp e_{u'v}^{(l,r,q)}}, \qquad (5)$$

$$m_v^{(l,r)} = \|_{q=1}^{H} \sum_{u \in N_r(v)} \alpha_{uv}^{(l,r,q)} \tilde{h}_u^{(l,r,q)} + c^{(l,r)} \in R^{Hd}, \qquad (6)$$

where $\|$ denotes concatenation, $d_0 = n_{max}$ and $d_1 = Hd$. Messages are summed over the relations that end at the node's type $\tau(v)$.

$$h^{-(l)}_{v} = \sum_{r:target(r)=\tau(v)} m_v^{(l,r)}, \qquad (7)$$

So in the paired model TFs receive messages from REs, REs from TFs and TGs, and TGs from REs, and in the unpaired model TFs and TGs receive messages from each other. Each relation has its own parameters, so messages along different relations into the same node type are weighted separately.

**Normalization, residual and dropout.**

$$h_v^{(1)} = Drop\left(BN_\tau^{(1)}(\bar{h}^{(1)})_v\right), \qquad (8)$$

$$z_v^{(t)} = Drop\left(PN(BN_\tau^{(2)}(\bar{h}^{(2)}))_v + W_\tau^{res} x_v^{(t)} + b_\tau^{res}\right) \in R^{Hd}, \qquad (9)$$

where $BN_\tau$ is batch normalization over the nodes of type $\tau$ in one stage graph and $Drop$ is dropout. PairNorm centres the representations of a node type and rescales them, which keeps the representations of different nodes apart after message passing. The paired models scale each row to unit length,

$$PN(U)_v = \frac{U_v - \bar{U}}{\|U_v - \bar{U}\|_2}, \qquad \bar{U} = \frac{1}{N_\tau}\sum_w U_w, \qquad (10)$$

and the mouse model divides all rows by their mean length. Apart from attention, the encoder has no pointwise nonlinearity. The values of $H$, $d$ and the dropout rate are given under Optimization (Section 4).

#### 3 Temporal module[2]

**Carried recurrent state.** For each node type, a gated recurrent unit links the stages along the lineage tree. With input $z = z_v^{(t)}$ (equation (9)) and previous state $s = s_v^{(pa(t))}$,

$$\gamma^r = \sigma(W_r z + b_r + U_r s + c_r), \qquad \gamma^u = \sigma(W_u z + b_u + U_u s + c_u), \qquad (11)$$

$$\tilde{s} = tanh(W_n z + b_n + \gamma^r \odot (U_n s + c_n)), \qquad GRU_\tau(z, s) = (1 - \gamma^u) \odot \tilde{s} + \gamma^u \odot s, \qquad (12)$$

with reset gate $\gamma^r$, update gate $\gamma^u$, candidate state $\tilde{s}$, $\sigma$ the logistic function and state dimension $d$. The state carried to the child stages includes a linear skip connection of the current representation,

$$s_v^{(t)} = GRU_\tau\left(z_v^{(t)}, s_v^{(pa(t))}\right) + W_\tau^{skip} z_v^{(t)} + b_\tau^{skip}, \qquad s_v^{(pa(t))} = 0 \text{ at the root}, \qquad (13)$$

and the embedding is

$$y_v^{(t)} = W_\tau^{out} s_v^{(t)} + b_\tau^{out} \in R^d. \qquad (14)$$

The state thus mixes the current stage, through the gates and the skip term, with the history along the path from the root. Stages are processed in topological order, so the state of a branching stage is passed unchanged to each of its children; every root-to-leaf path is an unrolled sequence with the same parameters. A stage before a branch point is computed once and shared by the branches, not duplicated, and every stage enters the reconstruction loss once. $Y_{\tau}^{(t)}$ denotes the matrix whose rows are $y_v^{(t)}$, $v \in V_{\tau}$.

**Ablated variants (simulation).** Shared GAT omits the recurrent unit, $s_v^{(t)} = W^{skip} z_v^{(t)} + b^{skip}$, so each stage is embedded independently with shared parameters. LR+GRU replaces the two attention layers by node-type-specific linear maps, $\tilde{h}_v^{(1)} = W_{\tau}^{(1)} x_v^{(t)} + b_{\tau}^{(1)}$ and $\tilde{h}_v^{(2)} = W_{\tau}^{(2)} h_v^{(1)} + b_{\tau}^{(2)}$, and keeps everything else, so the prior graph enters only through the loss. Shared LR combines both changes. All four models use the same loss, sampler, inputs and number of epochs.

#### 4 Training objective

**Reconstruction of the prior.** DRAFT is trained to reconstruct the prior edges from the embeddings. For a prior relation $\varrho$ and stage $t$, the pair score is

$$p_{uv}^{(t)} = \sigma\left(\lambda_{uv}^{(t)}\right), \quad \lambda_{uv}^{(t)} = \langle y_u^{(t)}, y_v^{(t)} \rangle \text{ (paired)}, \qquad p_{uv}^{(t)} = \left|\cos\left(y_u^{(t)}, y_v^{(t)}\right)\right| \text{ (unpaired)}. \qquad (15)$$

In the paired model, the positives $P_{\varrho} = \{(u, v): A_{uv}^{\varrho} = 1\}$ are the same at every stage. At every epoch and stage, the same number of negatives $Q_{\varrho}^{(t)}$ is drawn uniformly without replacement from the non-edges $\{(u, v): A_{uv}^{\varrho} = 0\}$ of the relation. The loss of one relation and stage is the binary cross-entropy

$$L_{\varrho}^{(t)} = -\frac{1}{|P_{\varrho}| + |Q_{\varrho}^{(t)}|}\left(\sum_{(u,v)\in P_{\varrho}} \log p_{uv}^{(t)} + \sum_{(u,v)\in Q_{\varrho}^{(t)}} \log\left(1 - p_{uv}^{(t)}\right)\right), \qquad (16)$$

and $L_{rec} = \sum_t \sum_{\varrho} L_{\varrho}^{(t)}$ (the mean over stages and relations in the simulation). Because the negatives are redrawn at every epoch, each prior edge is contrasted with random non-edges of the same relation rather than with a fixed set. In the paired model TF–TG pairs are not a training target: TF and TG embeddings are coupled only through their shared RE neighbours.

**Temporal smoothness penalty.** Let $S_{\varrho}^{(t)}$ be the matrix of pair scores of relation $\varrho$ at stage $t$ over all node pairs, not only prior edges. The penalty measures how much these matrices change from a stage to its parent,

$$L_{smooth} = \sum_{b} w_b \frac{1}{|T_b^*||R_O|} \sum_{t\in T_b^*} \sum_{\varrho\in R_O} \nu_{\varrho} \| S_{\varrho}^{(t)} - S_{\varrho}^{(pa(t))} \|_F^2, \qquad (17)$$

where each $T_b^*$ is a set of stages that have a parent, $w_b$ is its weight and $\nu_\varrho$ a normalizing constant. The total loss is $L = L_{rec} + \omega\, L_{smooth}$, with $\omega$ set at the first epoch so that $\omega L_{smooth} = L_{rec}$ and then held fixed, so the two terms are balanced only at the start of training. The three implementations are special cases:

- *Drosophila*: one set, all stages with a parent ($w = 1$, $\nu_\varrho = 1$), and the logit matrices $S_{TR}^{(t)} = Y_{TF}^{(t)} Y_{RE}^{(t)\top}$ and $S_{RG}^{(t)} = Y_{RE}^{(t)} Y_{TG}^{(t)\top}$. Each term is computed from $d \times d$ Gram matrices,

  $$\|K_1 K_2^\top - K_3 K_4^\top\|_F^2 = \langle K_1^\top K_1, K_2^\top K_2\rangle_F + \langle K_3^\top K_3, K_4^\top K_4\rangle_F - 2\langle K_1^\top K_3, K_2^\top K_4\rangle_F, \quad (18)$$

  without forming the $N_{TF} \times N_{RE}$ or $N_{RE} \times N_{TG}$ matrices.

- Mouse: $S^{(t)}$ is the matrix of $|cos|$ scores of equation (15), $\nu = 1/(N_{TF} N_{TG})$, and the two sets are the stages of the successful and the unsuccessful branch, which share the step C4–C8.

- Simulation: no penalty ($\omega = 0$).

The penalty acts on the scores of the trained relations: TF–RE and RE–TG logits in *Drosophila* and the TF–TG score in the mouse.

**Scores after training.** After training, a TF–TG pair is scored at stage $t$ by $\hat{S}_{fg}^{(t)} = cos\left(y_f^{(t)}, y_g^{(t)}\right)$ in the paired model and by $p_{fg}^{(t)}$ of equation (15) in the mouse, among the active TFs and TGs $V_{TF}^{(t)}$ and $V_{TG}^{(t)}$, which are non-zero in at least 5% (*Drosophila*) or 30% (mouse) of the cells of the stage. The network of stage $t$ consists of its $k$ highest-scoring pairs, with $k = 10{,}000$ in *Drosophila* and $k = 50{,}000$ in the mouse.

**Optimization.** DRAFT and its internal ablations were trained full-batch with Adam across all stages jointly. Hyperparameters of paired and unpaired model and optimization settings are reported in the **Supplementary Tables 10 and 11**. Neither experimental reference networks nor simulated ground-truth networks were used for training or checkpoint selection.

#### 5 Evaluation metrics

This section defines the scores and metrics shared by all benchmarks; Sections 6–8 give, for each dataset, the design or preprocessing and the baselines, and for the mouse and *Drosophila* the evaluation protocol. The scripts that run each baseline with its arguments are in the code repository.

#### 5.1 Scores and ranking metrics

**DRAFT scores.** DRAFT is evaluated with the score of the relation that a reference covers: the TF–TG logit $\lambda_{fg}^{(t)}$ of equation (15) in the simulation, the RE–TG score $\sigma(\lambda_{jg}^{(t)})$ against *Drosophila* Hi-C, and the stored TF–TG score $p_{fg}^{(t)}$ (equation (15)) in the mouse. Baselines are ranked by their native edge scores, and pairs absent from a method's output score 0.

**Ranking metrics.** For evaluated pairs with labels $y \in \{0,1\}$ and scores $s$, let $TPR(\vartheta)$, $FPR(\vartheta)$ and $Prec(\vartheta)$ be the true-positive rate, false-positive rate and precision of the pairs with $s \geq \vartheta$. AUROC is the area under TPR against FPR as $\vartheta$ sweeps all score values; equivalently,

$$AUROC = \frac{1}{n_+ n_-} \sum_{a:y_a=1} \sum_{b:y_b=0} \left( 1[s_a > s_b] + \frac{1}{2} 1[s_a = s_b] \right), \qquad (19)$$

the probability that a random positive outranks a random negative, 0.5 for a random ranking. Average precision weights the precision at each distinct threshold $\vartheta_1 > \vartheta_2 > \ldots$ by the gain in recall, $AP = \sum_k (TPR(\vartheta_k) - TPR(\vartheta_{k-1}))Prec(\vartheta_k)$ with $TPR(\vartheta_0) = 0$. The mouse AUPR is the area under the precision–recall curve computed with the AUCCalculator of the original study[3], which interpolates between operating points. Both precision-based measures equal the positive fraction for a random ranking. All three depend only on the ranking of the scores, so no threshold has to be chosen.

#### 5.2 Temporal smoothness

Let $W_t$ be a method's score matrix at stage $t = 1, \ldots, T$ on a set of candidate pairs that is the same at every stage (pairs absent from the output score 0). With the unsquared Frobenius distance $\|W_i - W_j\|_F$ between stages,

$$C = 1 - \frac{\frac{1}{T-1}\sum_{t=1}^{T-1} \|W_{t+1} - W_t\|_F}{\max\limits_{i,j} \|W_i - W_j\|_F}. \qquad (20)$$

When all stage-specific networks are identical, temporal smoothness is defined as 1. $C \in [0,1]$ compares the mean step between adjacent stages with the largest change between any two stages. It is 1 if all $W_t$ are equal and does not depend on the scale of the scores, because each method is normalized by its own largest change. For DRAFT and its ablations in the simulation, $W_t$ holds the probabilities $\sigma(\lambda_{fg}^{(t)})$ on the candidate pairs; for the mouse, $C$ is computed separately for each branch. Heatmaps show the distances $\|W_i - W_j\|_F$ between all pairs of stages, divided by the largest where stated. $C$ measures how gradually the predicted networks change, not their accuracy: a time-invariant prediction attains $C = 1$, and a prediction that changes in one jump and is then constant can also score high.

#### 6 Simulation

##### 6.1 Design

**Modules and candidate regulators.** The simulated genome has $N_{TF} = 50$ TFs and $N_{TG} = 150$ TGs, each assigned at random to one of three modules of balanced size. For each TG $g$, nine candidate TFs $I_g$ are drawn from its own module and three, $O_g$, from the other modules. The candidate matrix is $M_{fg} = 1[f \in I_g \cup O_g]$, and the prior is modular because most candidates of a TG come from its own module.

**Dynamic networks.** At the first stage six TFs of $I_g$ are active, $A_g^{(0)}$; outside candidates are never active. At each transition $t-1 \to t$, $\Delta = round(0.1 \times 150 \times 6) = 90$ replacements are made, each exchanging, within one TG, a TF of $A_g^{(t-1)} \cap A_g^{(0)}$ for a TF of $I_g \setminus (A_g^{(t-1)} \cup A_g^{(0)})$, with the replacements drawn uniformly without replacement from all available exchange slots, $min\left(|A_g^{(t-1)} \cap A_g^{(0)}|, |I_g \setminus (A_g^{(t-1)} \cup A_g^{(0)})|\right)$ per TG. Every TG therefore keeps six active regulators, a removed edge never returns and an added edge is never removed. The ground-truth network is $A_{fg}^{*(t)} = 1[f \in A_g^{(t)}]$. Because the edges changed at different transitions are distinct,

$$\| A^{*(t)} - A^{*(t')} \|_F^2 = 2\Delta\,|t - t'|, \qquad (21)$$

so the distance between stages grows as $\sqrt{|t - t'|}$, and the temporal smoothness of the ground truth (equation (20) with $W_t = A^{*(t)}$) is $1 - 1/\sqrt{T-1} = 0.5$ for $T = 5$ stages.

**Regulatory elements and accessibility.** The 12 candidates of each TG are split at random into six pairs. Each pair $F(j)$ defines one RE $j$ that targets only that TG, so $A_{fj}^{TR} = 1[f \in F(j)]$, $A_{jg}^{RG} = 1[g \in G(j)]$ with $G(j) = \{g\}$, and equation (1) recovers $M$ exactly. These maps do not change over time. The activity of RE $j$ at stage $t$ is the fraction of its TFs that regulate its target,

$$a_j^{(t)} = \frac{|F(j) \cap A_g^{(t)}|}{|F(j)|} \in \left\{0, \frac{1}{2}, 1\right\}, \qquad (22)$$

and the latent accessibility of RE $j$ in the cells of stage $t$ is multiplied by $0.2 + 0.8\,a_j^{(t)}$.

**Expression and observation.** Expression and accessibility were simulated with scMultiSim[4] 1.2.0 along a linear tree of five segments (200 cells each), with the generating network of every cell set to $A^{*(t)}$ of its segment and native kinetic RNA simulation. Observed counts are independent Poisson draws, $o_{vc} \sim Poisson(\delta\, \tilde{o}_{vc})$ with depth $\delta = 1$, from the true counts $\tilde{o}_{vc}$ of RNA and ATAC. All methods receive the observed counts and the prior ($A^{TR}$ and $A^{RG}$, or $M$); DRAFT, SCENIC+ and scMTNI use $log(1 + x)$ of counts per $10^4$, and Dictys and scDoRI take the raw counts and normalize them themselves. $A^{*(t)}$, module labels and replacement events are used only for evaluation. Three datasets were generated (data seeds 101–103). In the branching design a shared trunk T0–T1 splits into two branches, $pa(A2) = pa(B2) = T1$, whose transitions are drawn independently with the rule above. Sensitivity settings and evaluation schemes are detailed in **Fig. 2c,d** and **Supplementary Fig. 3**.

**Prior SNR.** The modularity of a candidate graph is measured on its degree-normalized candidate matrix[5],

$$B = D_{TG}^{-1/2} M^{\top} D_{TF}^{-1/2}, \qquad SNR = \frac{\sigma_2(B)}{\sigma^*}, \qquad (23)$$

where $D_{TG}$ and $D_{TF}$ hold the row and column sums of $M^{\top}$, $\sigma_2(B)$ is the second singular value of $B$, and $\sigma^*$ is the largest $\sigma_2$ among three randomizations of $M$ that keeps every row and column sum (curveball algorithm). The first singular value of $B$ is always 1 and reflects the degrees alone; the second measures the strongest block structure, so $SNR \approx 1$ means no structure beyond the node degrees. For *Drosophila*, A links a TF to a TG when they share an RE in the prior (top 3% TF–RE motif pairs; RE–TG within 100 kb), restricted to measured genes and peaks and to the filtered TF list (208 TFs with candidates, 23,075 TGs). For mouse reprogramming, A is the union of the seven cluster-specific TF–TG priors used for training (score > 0.1). The ceiling is the largest second singular value of B over three degree-preserving Curveball shuffles of A. This gives SNR = 1.23 for *Drosophila* and 1.64 for the mouse union, 1.85, 1.67, 1.36 and 1.19 for the simulated priors with two, three, six and eight cross-module candidates **(Fig. 2c,d)**.

##### 6.2 Baselines

SCENIC+[6] is represented by its TF-to-gene module: gradient-boosting importances of all 50 TFs for each gene at each stage, without a prior. scMTNI[3] treats the stages as a lineage chain and uses the prior-connected TF–TG pairs as a soft prior; its edge scores are selection frequencies over 50 subsamples of half of the cells, with sparsity parameter $\beta_0 =- 3$. Dictys[7] fits its Ornstein–Uhlenbeck model at each stage with the prior-connected pairs as the allowed edges and scores an edge by its absolute normalized weight. scDoRI[8] is trained on the paired counts with the TF–RE prior (unsigned, in both its activator and repressor branches) and the RE–TG prior; the network of a stage is the cell-averaged mixture of its topic-specific TF–TG networks, in absolute value. Random scores are drawn from $Uniform(0,1)$.

#### 7 Mouse reprogramming

##### 7.1 Model inputs

The per-cluster model inputs (expression tables, regulator and gene lists, prior networks) and the cluster tree were generated with the processing code of the original study[3] and are identical to the processed files released with scMTNI (Zenodo 7879228). In this processing, ATAC fragments were counted in 1-kb bins and summed per gene with the nearest TSS, RNA was library-size normalized and log-transformed, the genes shared by both modalities were integrated with LIGER[9] ($k = 8$ factors) to define the clusters, and the lineage is the minimum spanning tree over cluster pseudobulk expression profiles.

##### 7.2 Baselines

SCENIC[10] infers co-expression modules with GRNBoost2 and prunes them with cisTarget motif enrichment; an edge is scored by its largest importance across the retained regulons. Dictys is run as in the simulation, with the DRAFT prior as the allowed edges and its expression covariance rank capped in the three smallest clusters. For scMTNI, the published scMTNI+Prior networks of the original study were used without refitting. Random networks are uniform TF–TG edge sets of the size of each cluster's prior.

##### 7.3 Evaluation

For cluster $t$ and gold standard $S$, the evaluated positives are $S_t = \{(f, g) \in S: f \in R_t,\ g \in X_t,\ f \neq g\}$, with $R_t$ and $X_t$ the regulator and gene lists of the cluster. With $S_t^{TF}$ and $S_t^{TG}$ the TFs and TGs that occur in $S_t$, the evaluated universe is

$$\Omega_t = \{(f, g) \in S_t^{TF} \times S_t^{TG}: f \neq g\}, \qquad (24)$$

all pairs between covered TFs and covered TGs, without restriction to prior edges. AUPR is computed on all of $\Omega_t$ at its natural prevalence $|S_t|/|\Omega_t|$ with the original study's preparation script. For AUROC, all positives and an equal number of negatives drawn uniformly from $\Omega_t \setminus S_t$ are split in half within each class, and AUROC is computed on one fixed half, the same pairs for every method. Results are the mean and sample standard deviation over the seven clusters.

#### 8 *Drosophila* testis

##### 8.1 Preprocessing

Nuclei were filtered on detected genes, counts and mitochondrial fraction, and doublets removed; ATAC nuclei were filtered on fragment counts, fraction of reads in peaks, TSS enrichment and nucleosome signal. Peaks were called with MACS3[11] and represented by TF–IDF and truncated singular value decomposition (latent semantic indexing); RNA principal components and ATAC LSI components were each corrected across samples with Harmony[12]. The prior has 761 TF nodes, and TF–TG networks are restricted to the 272 TFs of the filtered list by FlyBase[13].

#### 8.2 Baselines

For SCENIC+ region-to-gene scores, accessibility was imputed from pycisTopic topics computed once across all cells, and the gradient-boosting regression of gene expression on region accessibility was fitted on the cells of each stage. scDoRI used the binary TF–RE support of the DRAFT prior in place of motif scanning, filtered by its TF–peak correlation step, with RE–TG weights initialized by a distance decay. The Mondo timing comparison **(Supplementary Fig. 10)** used the published SCENIC+ eRegulons of the source study[14].

#### 8.3 Hi-C evaluation[15]

At stage $t$, the evaluated pairs are the prior RE–TG edges shared by all methods. The positives $P_t^{HiC}$ are those with a loop (2-kb resolution) whose one anchor overlaps the TSS of the TG and whose other anchor overlaps the peak; SpG loops label Sp-2 to Sp-16 and SpC loops *1*° Sc_1 to *2*° Sc_6. An equal number of negatives is drawn uniformly with replacement from the prior pairs without loop support. The unique pairs of each class are split 50:50, with repeated negative draws kept in one half, and AUROC is computed on one fixed half. The same pairs are used at every stage of an assay and for every method, and each stage contributes one value. The random control draws, at each stage, n_prior distinct pairs from the full RE×TG product, n_prior being the number of prior RE–TG edges, and scores each evaluated pair by its membership.

#### 8.4 Cross-lineage alignment and ligand–receptor timing

**Notation and dense networks.** $\Lambda \in \{S, G\}$ denotes the somatic or germline lineage, with $T_S = 9$ and $T_G = 15$ stages $t = 1, \ldots, T_\Lambda$ numbered in the order given in the Methods (SdA and SdB are the last two somatic stages); the hub is a separate sender group. Expression values are log-normalized counts, and $\phi_v^{(t)}$ is the fraction of the cells of stage $t$ in which gene $v$ is detected. Each lineage is given a continuous time axis by resampling its stage embeddings on a grid $\xi_1, \ldots, \xi_{\tilde{T}_\Lambda}$ with four points between consecutive stages,

$$\kappa_t(\xi) = \frac{exp\left(-(\xi - t)^2/2h^2\right)}{\sum_{t'} \; exp\left(-(\xi - t')^2/2h^2\right)}, \qquad y_v(\xi) = \sum_t \; \kappa_t(\xi)\, y_v^{(t)}, \qquad h = 0.7, \qquad (25)$$

a weighted average of neighbouring stages, with the trained embedding used exactly at integer $\xi$. TF–TG pairs of the resulting dense networks are scored by the cosine similarity $\tilde{S}^{(\Lambda)}_{fg}(\xi)$ of the resampled embeddings, and a retained pair has weight $\pi^{(\Lambda)}_{fg}(\xi) = \left(\tilde{S}^{(\Lambda)}_{fg}(\xi) + 1\right)/2 \in [0,1]$.

**Network fingerprints.** At each grid point $\xi_s$, let $\widehat{A}^{(\Lambda)}_s$ be the weighted TF–TG adjacency of the retained pairs, restricted to the TFs and TGs present in both lineages. It is normalized symmetrically by degree,

$$\widehat{A}^{(\Lambda),norm}_s = diag(\widehat{A}^{(\Lambda)}_s 1)^{-\frac{1}{2}} \widehat{A}^{(\Lambda)}_s \, diag(\widehat{A}^{(\Lambda)\top}_s 1)^{-\frac{1}{2}}, \qquad (26)$$

so that its singular values describe the structure of the network rather than its size. The fingerprint $\sigma^{(\Lambda)}_s$ collects the 20 leading singular values, each z-scored across the grid points of the lineage. Time is normalized to $\tau^{(\Lambda)}_s = (\xi_s - 1)/(T_\Lambda - 1) \in [0,1]$.

**Dynamic time warping.** Somatic grid points $s$ are matched to germline grid points $s'$ by the cost

$$\Omega_{ss'} = \| \sigma^{(S)}_s - \sigma^{(G)}_{s'} \|^2_2. \qquad (27)$$

Dynamic time warping[16] finds the monotone path through the cumulative cost $\Psi_{ss'} = \Omega_{ss'} + min\{\Psi_{s-1,s'}, \Psi_{s,s'-1}, \Psi_{s-1,s'-1}\}$ within a band of $\pm 6$ grid points around the diagonal. Each somatic grid point takes the mean normalized germline time of the points it is matched to, $w^0_s$, with $w^0_1 = 0$ and $w^0_{\tilde{T}_S} = 1$, and the warp is smoothed by

$$\widehat{w} = \underset{w}{argmin} \; \| w - w^0 \|^2_2 + \eta \sum_s (w_{s+1} - 2w_s + w_{s-1})^2, \qquad w_1 = 0,\; w_{\tilde{T}_S} = 1, \qquad (28)$$

with $\eta = 1$; the result is clipped to $[0,1]$ and made non-decreasing by isotonic regression. A somatic grid point is placed at the germline stage coordinate $1 + \widehat{w}_s(T_G - 1)$. The path was computed once and stored. For the scores, SdA and SdB form one terminal somatic stage placed at the last germline stage, and the path runs linearly to it from ISCC-3. The linear baseline places the eight somatic stages at equal spacing over the germline stages.

**Anatomical anchor.** In each lineage the switch is the largest increase of a stage-mean profile between adjacent stages within the spermatogonia-to-spermatocyte region, placed at the midpoint of its step: the expression of *eya* in the soma and the mean of *kmg*, *aly* and *can*,[17] each scaled to $[0,1]$, in the germline. The expected window of the mapped somatic switch is from Sp-8 to the germline switch.[18] The support of a switch is the fraction of bootstrap resamples of cells in which the same step is the largest.

**Ligand–receptor scores.** For a ligand $\ell$ and a receptor $g \in V_{TG}$, scores are computed on the stages $t$ of the receiving lineage. A pair is a candidate in a direction if the ligand is detected in at least 5% of the cells of a sender group and the receptor in at least 5% of the cells of a receiver stage. The ligand term $L_\ell^{(t)}$ is the mean expression of $\ell$ in the sender group; it is constant for the hub, whose cells do not progress through stages,[19] and for somatic and germline senders it is read at the sender position aligned to receiver stage $t$. The receptor term uses the receiver's trained embeddings at the integer stages. With $\Pi^{(t)} = \{\hat{S}_{fg}^{(t)} : f \in V_{TF}^{(t)},\ g \in V_{TG}^{(t)}\}$ the cosine scores of all active pairs (Section 4), the percentile of receptor $g$ with TF $f$ is $\rho_{fg}^{(t)} = |\{x \in \Pi^{(t)} : x \leq \hat{S}_{fg}^{(t)}\}|/|\Pi^{(t)}|$, and the network centrality of the receptor is

$$R_g^{(t)} = \frac{1}{|V_{TF}^{(t)}|} \sum_{f \in V_{TF}^{(t)}} \rho_{fg}^{(t)}. \qquad (29)$$

The communication score is

$$Comm_{\ell g}^{(t)} = L_\ell^{(t)} R_g^{(t)}\, 1\left[\phi_g^{(t)} \geq 0.05\right], \qquad (30)$$

so a stage at which the receptor is detected in fewer than 5% of the receiver cells scores 0.

**Peak stage and earliness.** A pair is unscored if $Comm_{\ell g}^{(t)} = 0$ at every stage and untimed if its relative dynamic range, $(max_t Comm_{\ell g}^{(t)} - min_t Comm_{\ell g}^{(t)})/max_t Comm_{\ell g}^{(t)}$, is below 0.1. Otherwise its peak stage is early (germline GSC to Sp-16; somatic CySC to eSCC-3) or late, and the earliness score is

$$ES_{\ell g} = \frac{\max_{t \in early} Comm_{\ell g}^{(t)} - \max_{t \in late} Comm_{\ell g}^{(t)}}{\max_t Comm_{\ell g}^{(t)}} \in [-1, 1], \qquad (31)$$

and 0 for unscored pairs. The expression-based comparators (ligand–receptor expression product, CellChat[20] and LIANA+[21]) replace $L_\ell^{(t)} R_g^{(t)}$ by their own group-pair scores, read along the same alignment and gated by the same receptor detection; peak stages and earliness are then derived in the same way.

**Literature benchmark.** For each direction with more than one literature pair, the earliness scores of the literature pairs $P^{lit}$ are compared with those of random size-matched sets $Q_b$ of non-literature candidates,

$$\underline{AUROC} = \frac{1}{B} \sum_{b=1}^{B} AUROC\left(\{ES_p\}_{p \in P^{lit}}, \{ES_p\}_{p \in Q_b}\right), \qquad |Q_b| = |P^{lit}|, \qquad (32)$$

with the same draws for every method. Under a uniformly random peak stage, the expected fraction of literature pairs peaking early is the fraction of early stages among the receiver's stages.

#### References


1. Veličković, P. *et al.* Graph Attention Networks. Preprint at https://doi.org/10.48550/arXiv.1710.10903 (2018).
2. Cho, K. *et al.* Learning Phrase Representations using RNN Encoder–Decoder for Statistical Machine Translation. in *Proceedings of the 2014 Conference on Empirical Methods in Natural Language Processing (EMNLP)* (eds Moschitti, A., Pang, B. & Daelemans, W.) 1724–1734 (Association for Computational Linguistics, Doha, Qatar, 2014). doi:10.3115/v1/D14-1179.
3. Zhang, S. *et al.* Inference of cell type-specific gene regulatory networks on cell lineages from single cell omic datasets. *Nat. Commun.* **14**, 3064 (2023).
4. Li, H., Zhang, Z., Squires, M., Chen, X. & Zhang, X. scMultiSim: simulation of single-cell multi-omics and spatial data guided by gene regulatory networks and cell-cell interactions. *Nat. Methods* **22**, 982–993 (2025).
5. Dhillon, I. S. Co-clustering documents and words using bipartite spectral graph partitioning. in *Proceedings of the seventh ACM SIGKDD international conference on Knowledge discovery and data mining* 269–274 (Association for Computing Machinery, New York, NY, USA, 2001). doi:10.1145/502512.502550.
6. Bravo González-Blas, C. *et al.* SCENIC+: single-cell multiomic inference of enhancers and gene regulatory networks. *Nat. Methods* **20**, 1355–1367 (2023).
7. Wang, L. *et al.* Dictys: dynamic gene regulatory network dissects developmental continuum with single-cell multiomics. *Nat. Methods* **20**, 1368–1378 (2023).
8. Saraswat, M. *et al.* Decoding Plasticity Regulators and Transition Trajectories in Glioblastoma with Single-cell Multiomics. 2025.05.13.653733 Preprint at https://doi.org/10.1101/2025.05.13.653733 (2025).

9. Welch, J. D. *et al.* Single-Cell Multi-omic Integration Compares and Contrasts Features of Brain Cell Identity. *Cell* **177**, 1873-1887.e17 (2019).
10. Aibar, S. *et al.* SCENIC: single-cell regulatory network inference and clustering. *Nat. Methods* **14**, 1083–1086 (2017).
11. Doherty, P. *et al.* MACS3: A Peak-calling Platform for Bulk and Single-cell Regulatory Genomics. *Genomics Proteomics Bioinformatics* qzag097 (2026) doi:10.1093/gpbjnl/qzag097.
12. Korsunsky, I. *et al.* Fast, sensitive and accurate integration of single-cell data with Harmony. *Nat. Methods* **16**, 1289–1296 (2019).
13. Öztürk-Çolak, A. *et al.* FlyBase: updates to the Drosophila genes and genomes database. *Genetics* **227**, iyad211 (2024).
14. van Nierop y Sanchez, P. *et al.* Dissecting the enhancer gene regulatory network in early Drosophila spermatogenesis. *Nat. Commun.* **16**, 6766 (2025).
15. Ilyin, A. A. *et al.* Comparison of genome architecture at two stages of male germline cell differentiation in Drosophila. *Nucleic Acids Res.* **50**, 3203–3225 (2022).
16. Sakoe, H. & Chiba, S. Dynamic programming algorithm optimization for spoken word recognition. *IEEE Trans. Acoust. Speech Signal Process.* **26**, 43–49 (1978).
17. White-Cooper, H., Leroy, D., MacQueen, A. & Fuller, M. T. Transcription of meiotic cell cycle and terminal differentiation genes depends on a conserved chromatin associated protein, whose nuclear localisation is regulated. *Development* **127**, 5463–5473 (2000).
18. Fabrizio, J. J., Boyle, M. & DiNardo, S. A somatic role for eyes absent (eya) and sine oculis (so) in Drosophila spermatocyte development. *Dev. Biol.* **258**, 117–128 (2003).
19. Kiger, A. A., Jones, D. L., Schulz, C., Rogers, M. B. & Fuller, M. T. Stem Cell Self-Renewal Specified by JAK-STAT Activation in Response to a Support Cell Cue. *Science* **294**, 2542–2545 (2001).
20. Jin, S. *et al.* Inference and analysis of cell-cell communication using CellChat. *Nat. Commun.* **12**, 1088 (2021).

21. Dimitrov, D. *et al.* LIANA+ provides an all-in-one framework for cell–cell communication inference. *Nat. Cell Biol.* **26**, 1613–1622 (2024).

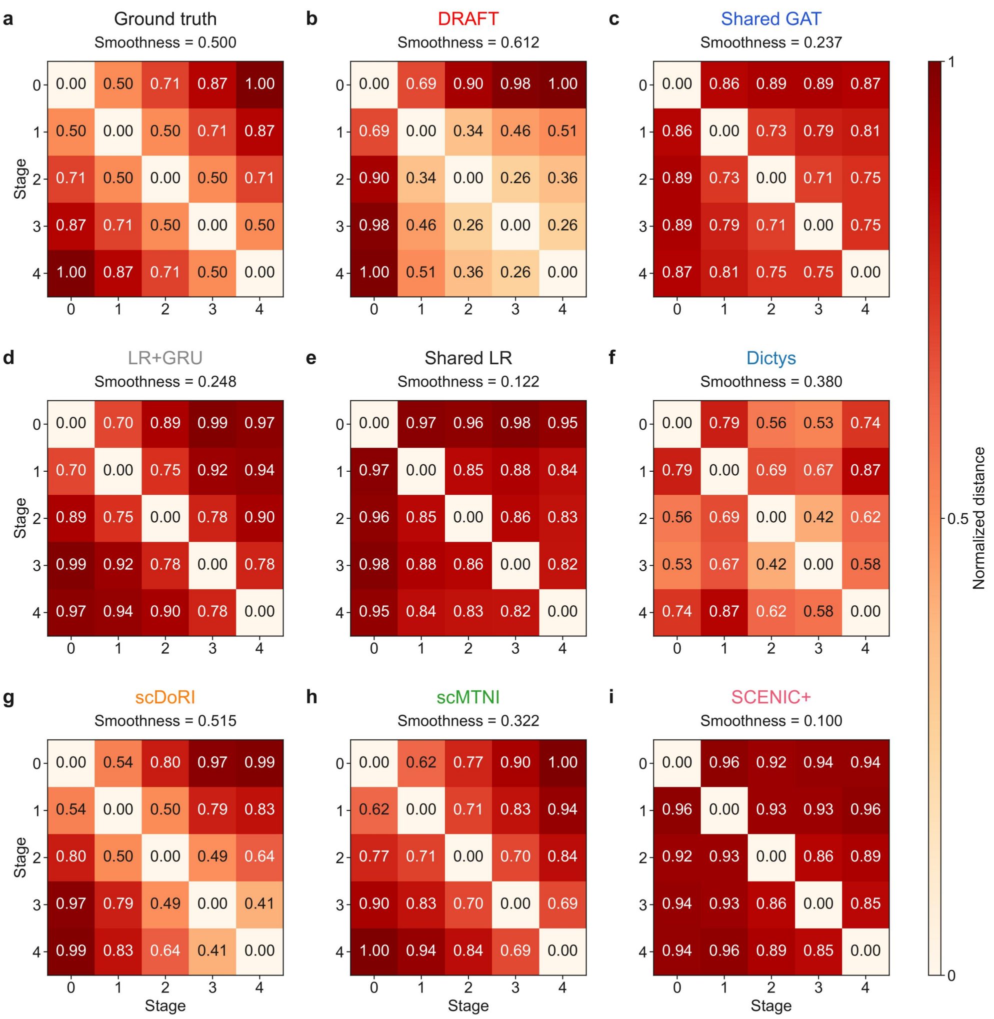


**Supplementary Fig. 1 | Pairwise network distances for the ground truth, DRAFT, its ablations and the benchmark methods in Fig. 2f.** Frobenius distances were computed over the same 1,800 candidate TF–TG pairs and divided by each run's maximum pairwise distance before averaging. Rows and columns represent stages T0–T4; titles give temporal smoothness **(Fig. 2f; Methods)**. Unnormalized maximum distances (mean ± s.d.) were 26.833 ± 0.000 for ground truth, 8.232 ± 0.309 for DRAFT, 4.344 ± 0.547 for Shared GAT, 2.212 ± 0.209 for LR+GRU, 2.132 ± 0.065 for Shared LR, 39.897 ± 25.491 for Dictys, 0.1065 ± 0.0084 for scDoRI, 10.293 ± 0.648 for scMTNI and 46.194 ± 0.589 for SCENIC+. Five model seeds were averaged within each of three independently simulated datasets, followed by averaging across datasets; s.d. is calculated across the three dataset means. Ground truth has one network sequence per dataset. Raw maxima retains method-specific score units and are not directly comparable across methods. Local fixed-input diagnostics suggest that DRAFT's initial jump partly reflects recurrent start-up, with increasing embedding norms shifting the overall magnitude of predicted edge scores.

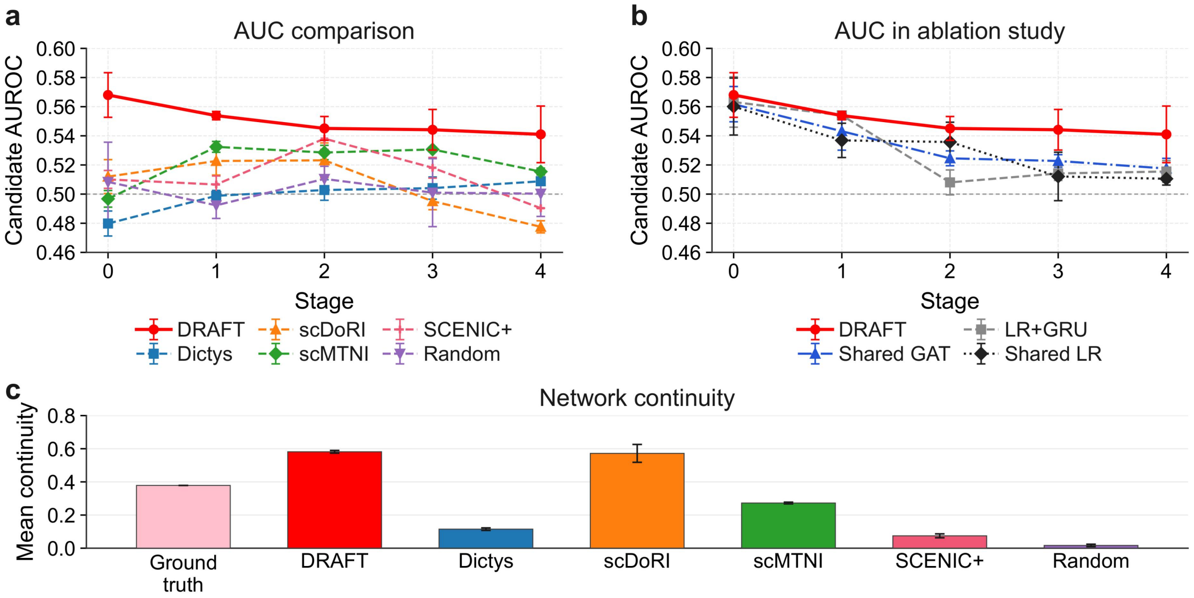


**Supplementary Fig. 2 |** Paired single-cell RNA-seq and ATAC-seq data were generated with scMultiSim for one fixed random-network realization comprising 50 TFs, 150 TGs and five stages, with 200 cells per stage. Each TG had 9 randomly sampled candidate TFs without module constraints; 6 were active and 3 inactive at each stage. Adjacent stages replaced 10% of active TF–TG edges. **a,** Candidate AUROC for DRAFT, SCENIC+, scMTNI, scDoRI and Dictys at each stage. Random denotes independent uniform edge scores. All methods were evaluated on the same TF–TG pairs in the prior network; dashed horizontal lines indicate chance AUROC (0.5). **b,** Ablation comparison of DRAFT, Shared GAT, LR+GRU and Shared LR. **c,** Mean temporal smoothness score evaluated on **Fig. 2**. Higher smoothness score indicates smoother score trajectories, without implying more accurate edge recovery.

Points and bars show means with sample standard deviations across three model initializations on the same dataset. The deterministic ground-truth reference has no initialization variability. In a separate control, shuffling edge scores within each scDoRI topic while retaining the same shuffled topic networks across time and the original mixing weights still yielded a mean smoothness of 0.611, indicating that high smoothness alone does not demonstrate recovery of true regulatory changes.

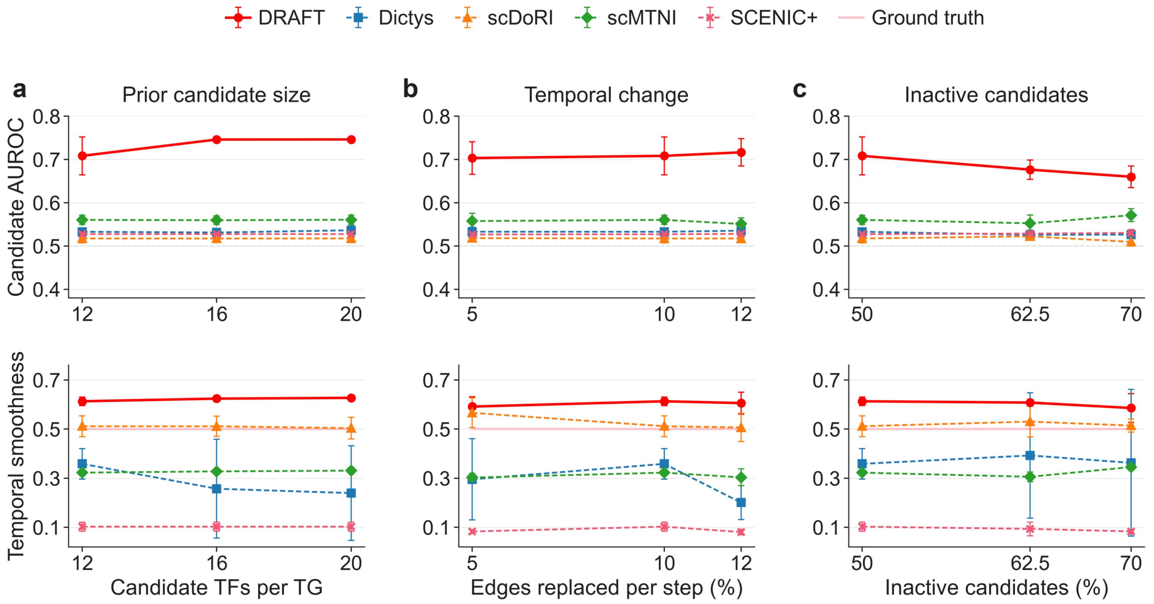


**Supplementary Fig. 3 | Sensitivity to candidate-prior size, temporal change and the proportion of inactive candidates.** Top row, candidate-set AUROC; bottom row, temporal smoothness **(Methods)**. **a,** Candidate-prior size was increased from 12 to 16 and 20 TFs per TG using nested priors. Ground-truth GRNs, RNA/ATAC observations, RE identities and RE–TG mappings were held fixed, while TF–RE links were expanded. All conditions were evaluated on the same 1,800 reference candidate pairs. **b,** The fraction of active edges replaced between adjacent stages varied across 5%, 10% and 12% (main design, 10%). **c,** The proportion of candidates inactive at each stage varied across 50%, 62.5% and 70%. For each condition, metrics were averaged over three model seeds within each of three independently simulated datasets. Points show the mean of the three dataset-level averages, and error bars indicate their sample standard deviation.

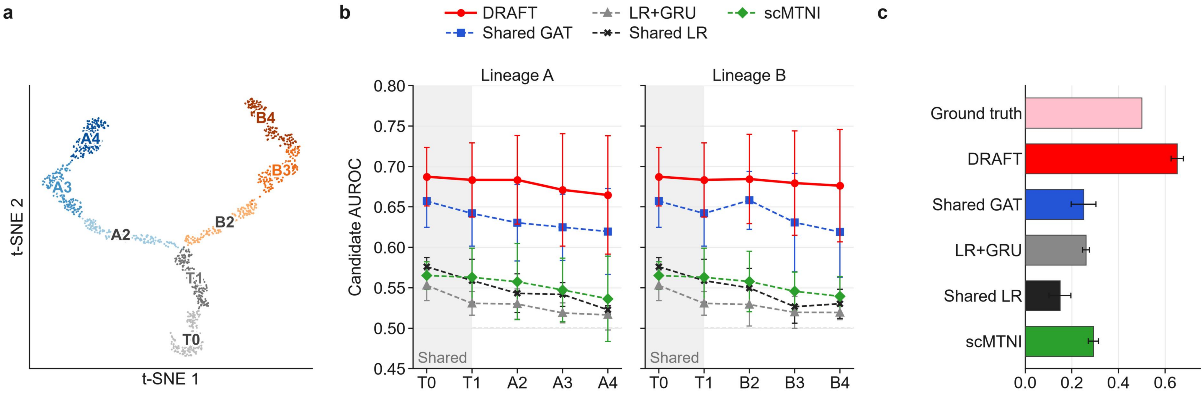


**Supplementary Fig. 4 | GRN inference on a branching simulated lineage.** The main simulation (Fig. 2) was extended to a lineage tree: two shared stages (T0, T1) followed by two branches of three stages each (A2–A4, B2–B4), with edge replacements drawn independently on each branch. All other settings, priors, training and scoring are as in the main simulation. DRAFT and its ablated variants received the tree through their parent–child relations, and scMTNI as its native lineage tree. SCENIC+, scDoRI and Dictys were not run, as they do not support branching lineages as applied in this benchmark. **a** RNA t-SNE of one dataset (1,600 cells); colors mark stages. **b** AUROC at each stage along lineage A (left) and B (right); T0 and T1 (shaded) are shared. **c** Temporal smoothness over the seven parent–child pairs of the tree; ground truth is the generating GRN. In **b** and **c**, values are means over three simulated datasets with three model seeds each; error bars, s.d. across datasets.

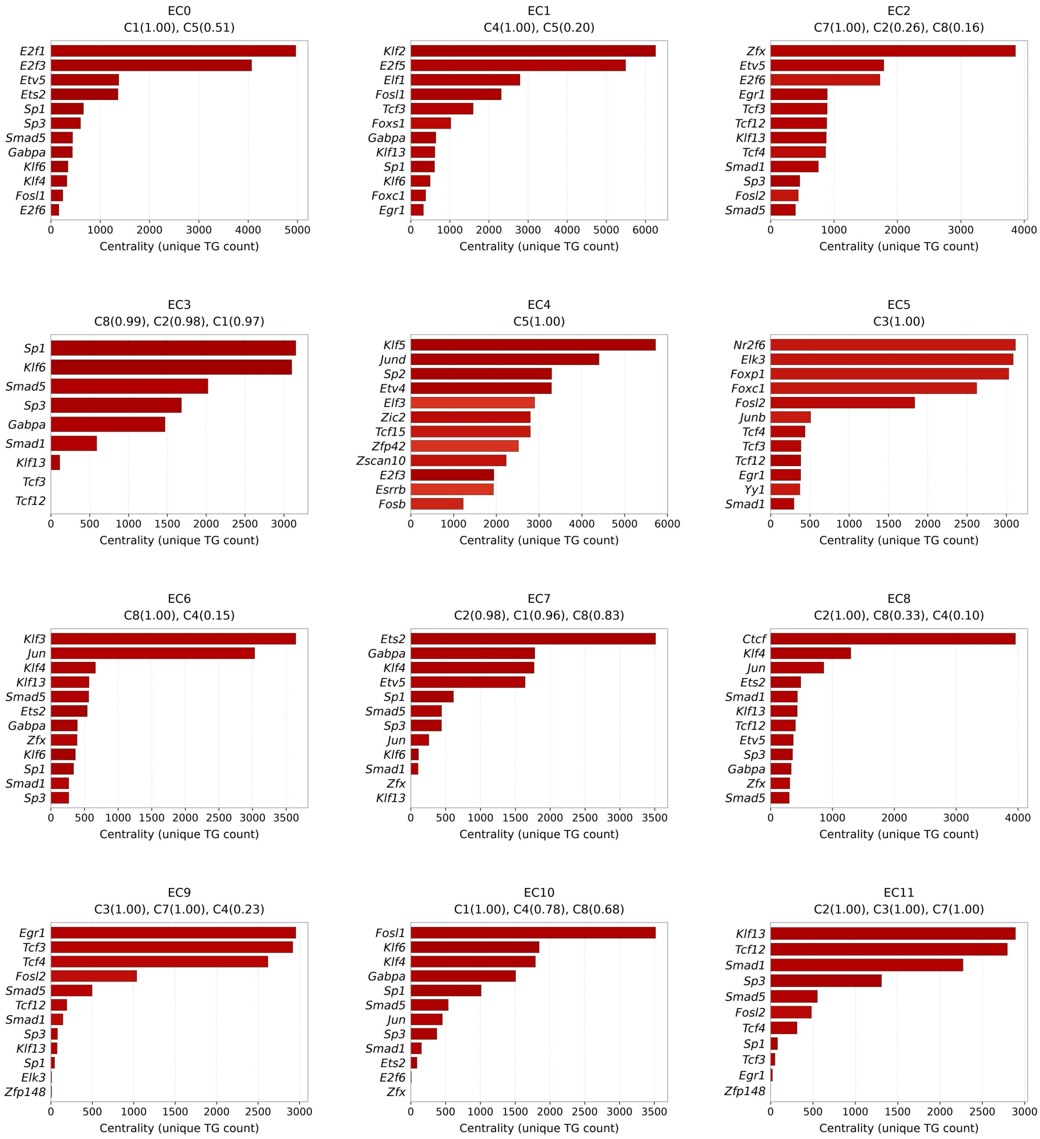


**Supplementary Fig. 5 |** Up to 12 transcription factors (TFs) with the highest centrality are shown for each of the 12 edge clusters (EC0–EC11) identified by k-means clustering of the reconstructed DRAFT networks. Bar length indicates the number of distinct inferred target genes connected to each TF within the corresponding EC, aggregated across cell clusters. Bar color reflects the mean inferred edge score for each TF–EC pair. Panel titles list up to three cell clusters with the highest representation of each EC (fraction ≥0.10). Values in parentheses denote the fraction of unique edges in that EC present in the indicated cell cluster. Because edges can occur in multiple cell clusters, these fractions need not sum to one. Centrality describes inferred network topology and does not directly measure TF activity, protein abundance or binding occupancy.

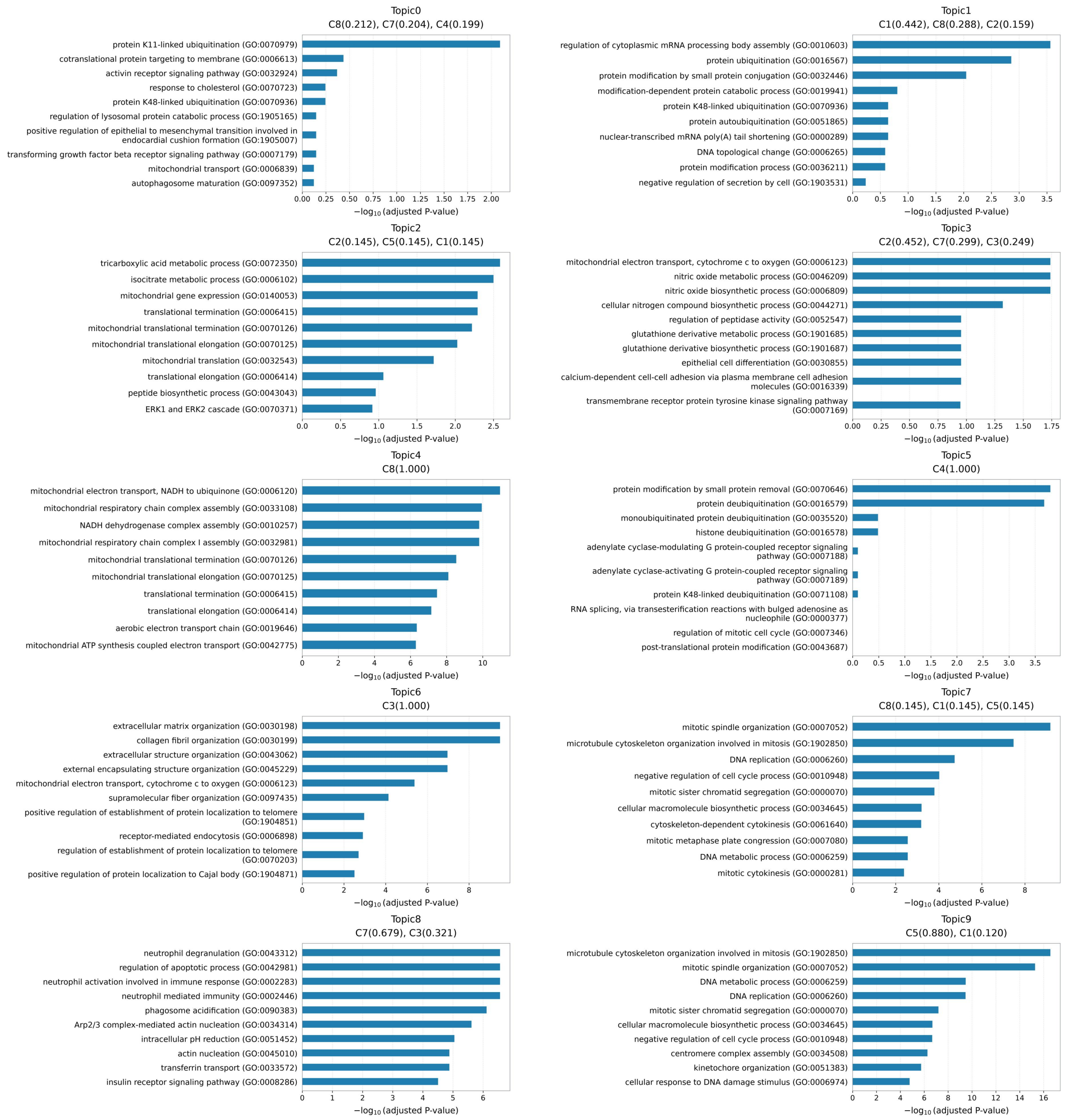


**Supplementary Fig. 6 |** Functional annotation of LDA-derived gene topics in mouse reprogramming. Gene Ontology (GO) Biological Process enrichment was performed for gene sets assigned to each of the ten topics (T0–T9) inferred from the reconstructed DRAFT networks. Analysis used the GO Biological Process 2021 collection and 7,764 network target genes as the background. P values were calculated using one-sided hypergeometric tests and adjusted using the Benjamini–Hochberg procedure within each topic. Bars show $-\log_{10}$(adjusted P value) for the ten highest-ranked terms per topic. Panel titles list up to three cell clusters with the largest normalized topic weights (weight ≥0.10), with weights indicated in parentheses. Weights are normalized across all seven cell clusters separately for each topic. Terms are displayed without an additional significance filter; therefore, not all displayed terms meet FDR <0.05.

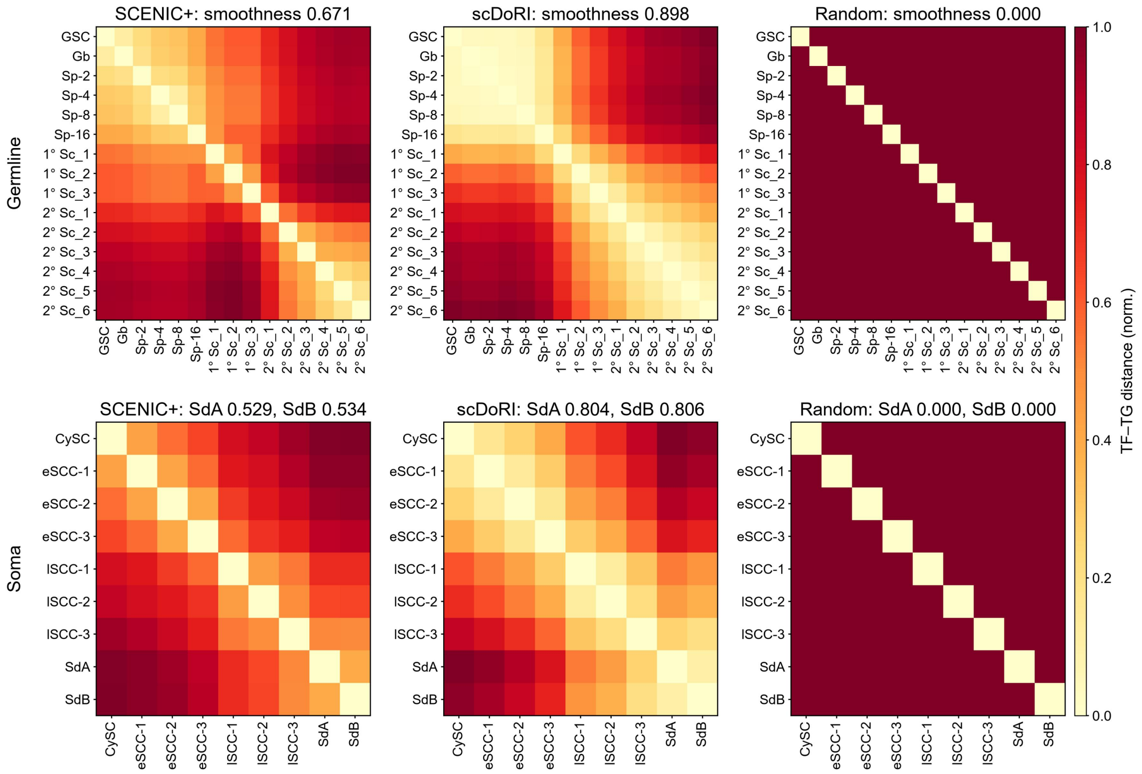


**Supplementary Fig. 7 | Temporal smoothness of baseline TF–TG networks.** Heatmaps show pairwise Frobenius distances between stage TF–TG networks for SCENIC+, scDoRI and random networks along the germline (top) and somatic (bottom) trajectories.

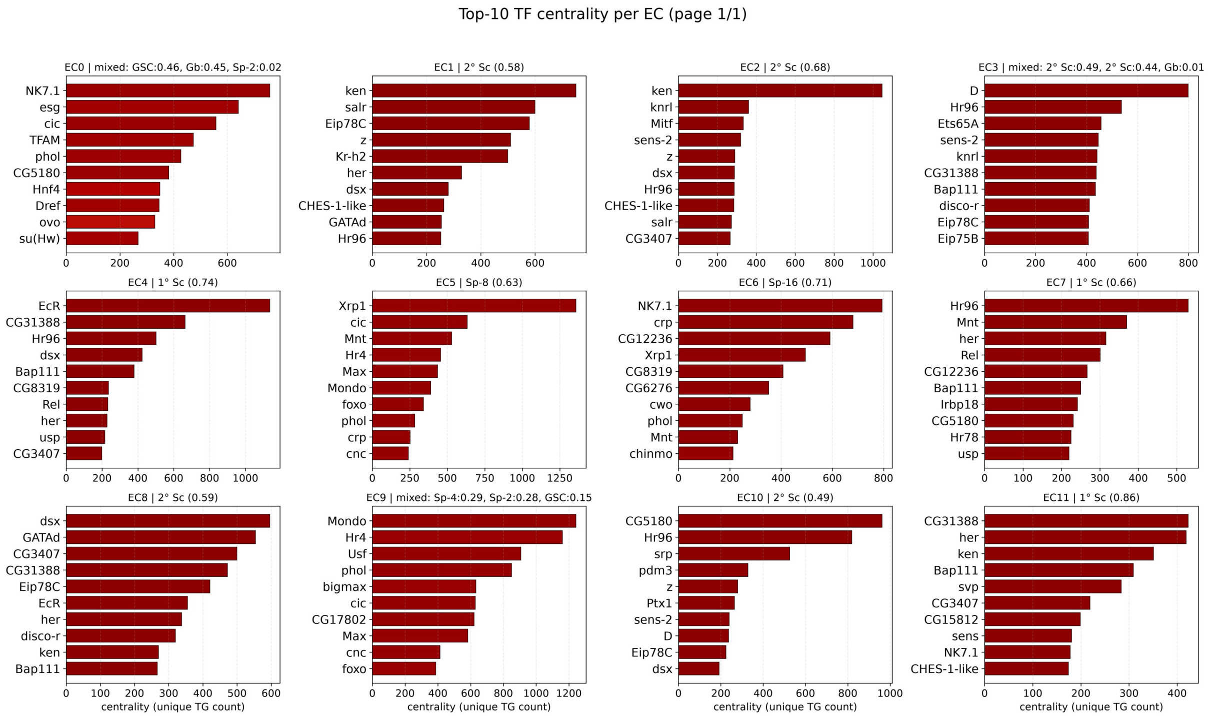


**Supplementary Fig. 8 |** The top 10 TFs ranked by centrality for each edge cluster. *Hr96*, which ranks first in EC7, has no reported testis function and is therefore not shown in **Fig. 4g**.

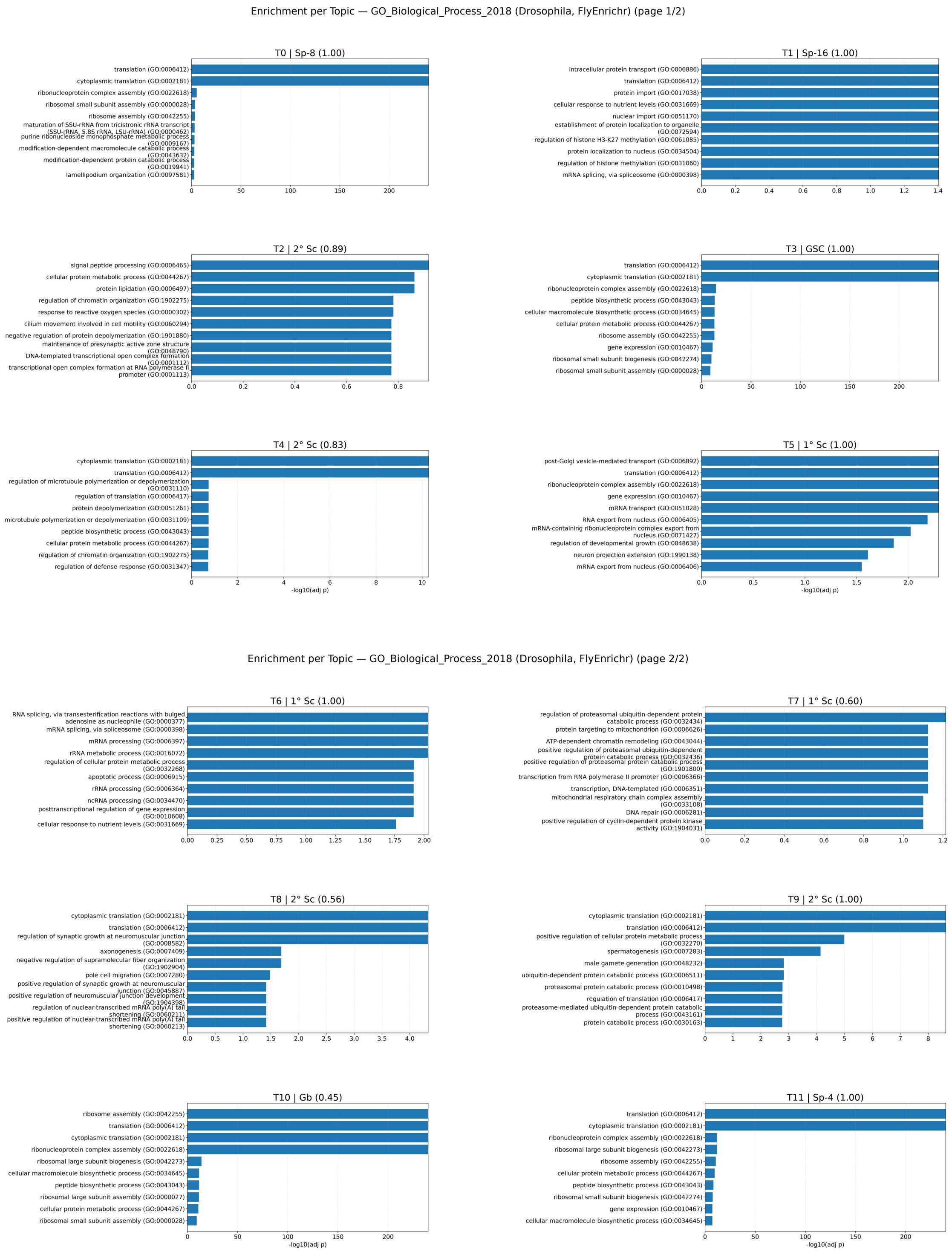


**Supplementary Fig. 9 |** The top active functional pathways identified via GO enrichment for each gene topic.

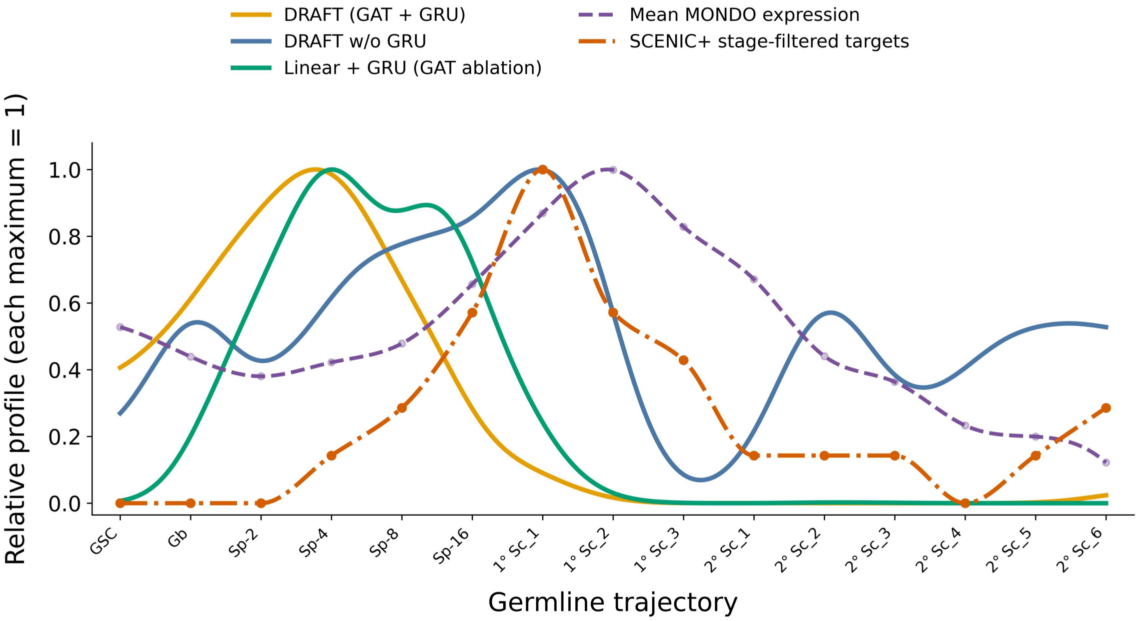


**Supplementary Fig. 10 |** Gold, blue and green curves show the network centrality of Mondo inferred by full DRAFT (GAT + GRU), DRAFT without GRU, and the GAT-to-Linear ablation retaining GRU, respectively (seed 2025 for all models). Centrality is the sum of Mondo's selected outgoing TF–TG edge scores at each dense stage, smoothed with a Gaussian kernel (σ = 0.5 stages). The purple dashed curve shows the stage-wise mean log-normalized Mondo expression, including zero-expression cells. The orange dash-dot curve shows the number of unique Mondo target genes reported by SCENIC+ in the original study, with genes linked to multiple regulatory regions counted once. Dots mark stage-level expression means and SCENIC+ counts; curves between them are cubic interpolations (shape-preserving for SCENIC+) for display only and are not additional observations. Because edge-scoring schemes differ between methods, each profile is normalized to its own maximum, so the panel compares temporal patterns rather than absolute expression, network strength or target counts. No scDoRI profile is shown because Mondo's candidate TF–RE connection was removed by correlation-based filtering during preprocessing, leaving no binding support for downstream network inference.

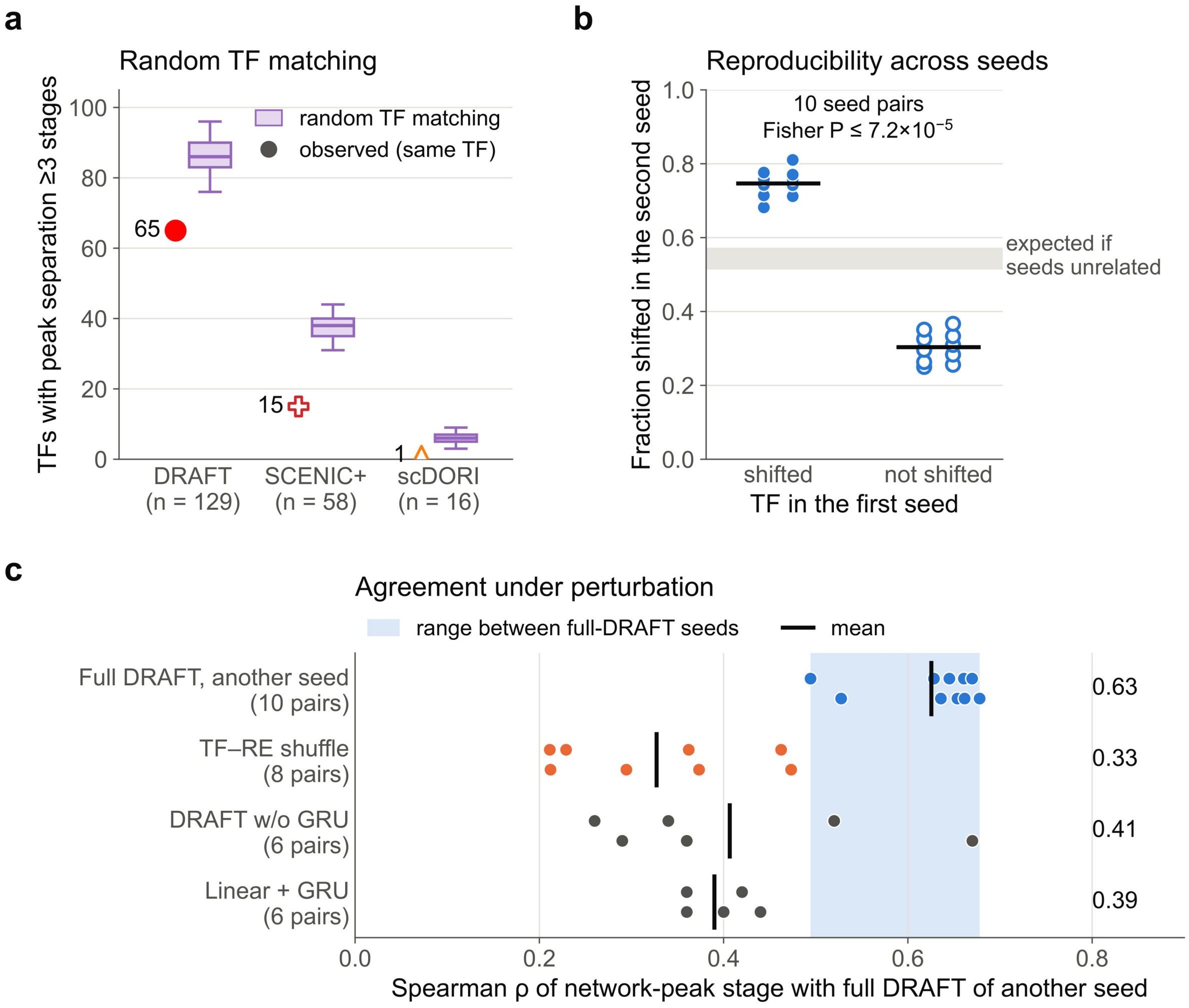


**Supplementary Fig. 11 | Controls for peak shifts between TF network activity and expression.** TF has a peak shift when its network peak and its expression peak lie ≥3 developmental stages apart (peaks: stages within 95% of the profile maximum). Network profiles are summed TF–TG scores (DRAFT), target counts (SCENIC+) or summed absolute TF–TG weights (scDoRI) per stage; expression profiles are mean RNA expression per stage. **a** Number of TFs with a peak shift in each method (symbols; n, evaluable TFs), compared with 10,000 random re-pairings of network and expression profiles among the same TFs (purple boxes: median, interquartile range and 2.5th–97.5th percentiles). Random pairs give at least as many shifts, so a shift alone does not indicate biological relevance. **b** Reproducibility across five training seeds (2024–2028; 2025 is the manuscript model). For each of the 10 seed pairs (dots), the fraction of TFs shifted in the second seed among TFs that were shifted (filled) or not shifted (open) in the first seed; black lines, means; grey band, the fraction expected if the two seeds were unrelated. $P \leq 7.2 \times 10^{-5}$ for every pair (one-sided Fisher's exact test). **c** Agreement of network-peak stages (Spearman ρ across TFs) between full DRAFT and models trained with a different seed: full DRAFT; DRAFT with a shuffled TF–RE prior (each RE keeps its number of TFs, which are drawn at random); DRAFT without the GRU; and Linear + GRU (GAT layers replaced by linear layers). Dots, model pairs (number in parentheses); black lines, means; blue band, range between full-DRAFT seeds.

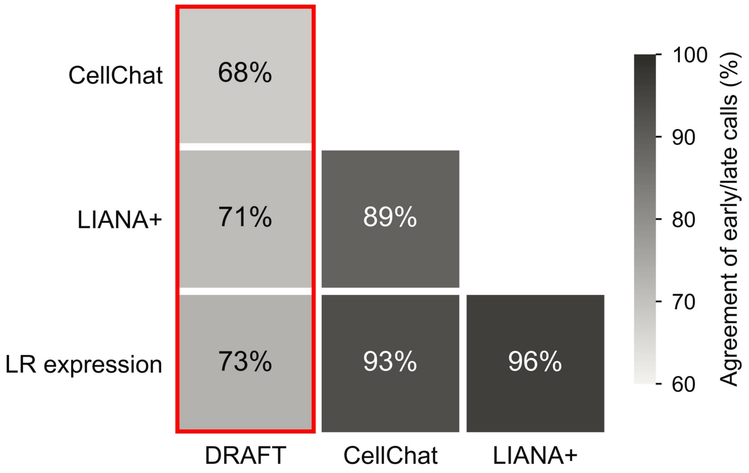


**Supplementary Fig. 12 | Agreement of early/late calls between methods.** Each cell gives the percentage of candidate LR pairs on which two methods make the same early/late call (Methods), counting only pairs timed by both methods. Candidates are pooled over the four directions (hub→germline, hub→soma, soma→germline and germline→soma; 322 pairs). Red outline, DRAFT. DRAFT values are means over five germline models trained with different random seeds (ranges 66–70%, 69–72% and 71–75%); somatic receivers come from a single model. CellChat, LIANA+ and the LR expression product do not depend on the seed. All methods use the same stage groups, expression gates and alignment.

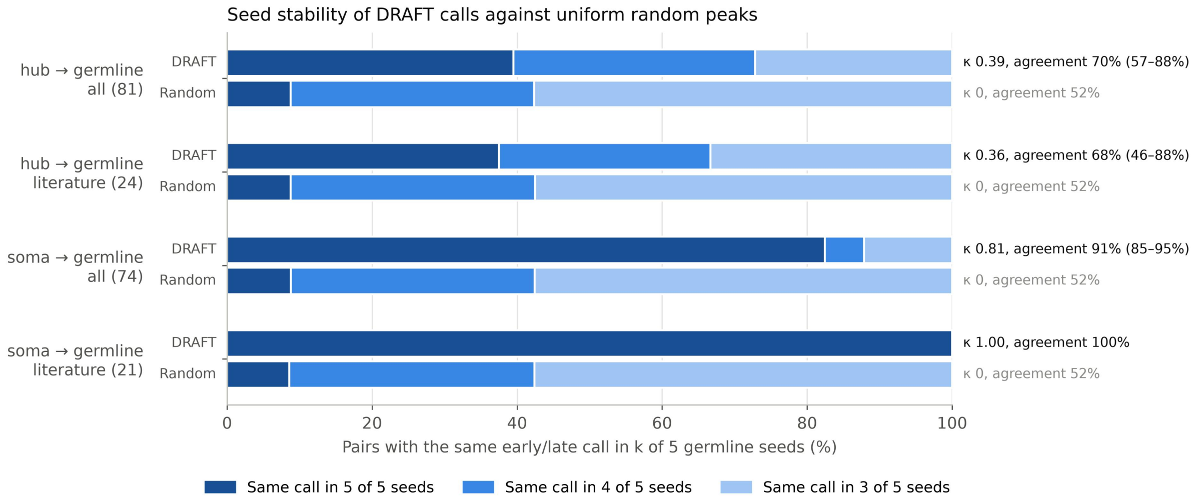


**Supplementary Fig. 13 | Stability of DRAFT early/late calls across germline models.** DRAFT receptor scores were computed with five germline models trained with different random seeds (2024–2028), and each candidate pair was called early or late in each model (Methods). Bars show the percentage of pairs that receive the same call in all five models, in four or in three. With five models and two classes, at least three models always agree. Rows show all candidate pairs and literature pairs for hub→germline and soma→germline; numbers of pairs are in parentheses. "Random" gives the proportions expected if each model placed each pair's peak on a uniformly random germline stage (early with probability 6/15). Right, Fleiss' κ over the five models and the agreement between two models, as the mean (range) over the ten model pairs. Ligand terms do not depend on the seed. Somatic receivers come from a single model and are not shown.

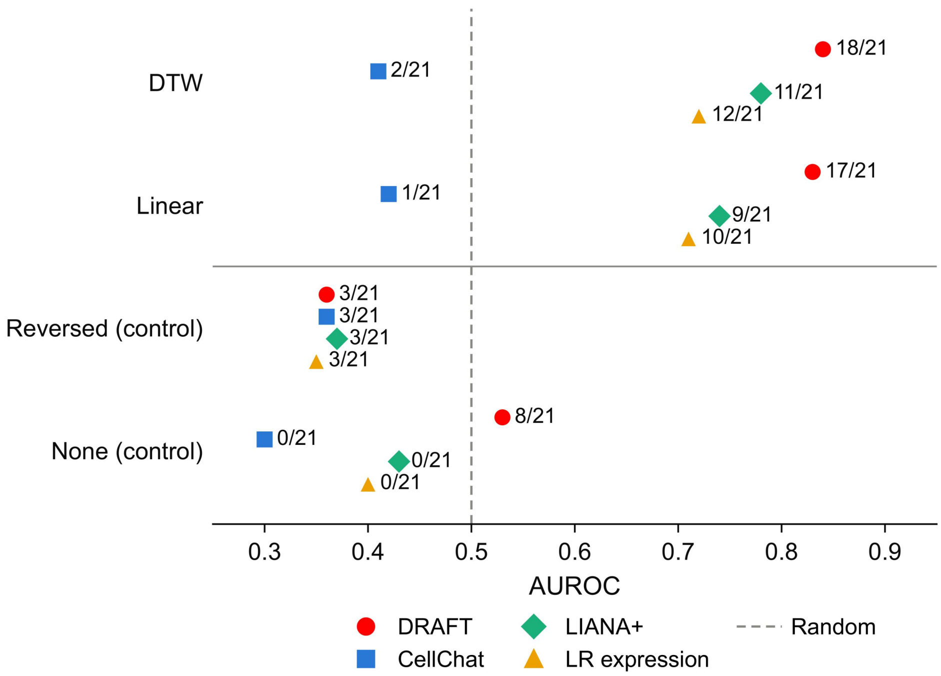


**Supplementary Fig. 14 | Soma→germline benchmark under different alignments.** AUROC of the earliness score **(Methods)** for the 21 literature-supported soma→germline pairs versus other candidate pairs. Each alignment determines which somatic stage is paired with each germline stage, and all four methods use the same alignment and expression gates. DTW, the alignment used in this study. Linear, constant developmental rate with the same start and end in both lineages. Reversed (control), linear alignment with somatic time reversed. None (control), somatic ligand expression averaged over all somatic groups. Labels, number of literature pairs whose score peaks in an early germline stage (GSC to Sp-16). Dashed line, random (AUROC 0.5). DRAFT scores come from the germline model trained with seed 2025.

**Supplementary Tables**

Supplementary Tables supplied as Excel workbooks

Supplementary Table 2 | Literature evidence for the transcription factors shown in **Fig. 4g**.

Supplementary Table 4 | TF network and RNA peak offsets.

Supplementary Table 7 | Cross-lineage alignment.

Supplementary Table 8 | Literature-supported ligand–receptor pairs.

Supplementary Table 9 | DRAFT early ligand–receptor candidates.

Supplementary Table 12 | Drosophila stocks.

Supplementary Table 13 | Antibodies and smFISH probes.

**Supplementary Table 1 | Key modeling differences between DRAFT and existing GRN inference methods.**

| Method | Input modalities | Temporal dependency modeling | Recurrent memory? | Nonlinear regulatory modeling? | Temporal constraint type | GRN includes RE? |
|---|---|---|---|---|---|---|
| DRAFT (paired) | RNA + ATAC | Lineage-aware GAT–GRU | Yes | Yes | Explicit constraint | Yes |
| DRAFT (unpaired) | Unpaired RNA + ATAC | Lineage-aware GAT–GRU | Yes | Yes | Explicit constraint | No |
| Dictys | RNA + ATAC | Trajectory-based moving windows | No | Stochastic model | Post-hoc smoothing/interpolation | No |
| CellOracle | RNA + ATAC priors | None | No | No | None | No |
| scMTNI | RNA + ATAC priors | Lineage-coupled multitask learning | No | Primarily linear | Explicit constraint | No |
| SCENIC+ | RNA + ATAC | None | No | Yes | None | Yes |
| scDoRI | RNA + ATAC | None | No | Yes | None | Yes |
| scTIE | RNA + ATAC | OT-based temporal alignment | No | Yes | Explicit constraint | Partial |
| Marlene | RNA | GRU-evolved attention parameters | Yes | Yes | Implicit coupling | No |

Methods are compared in how they handle dynamics and regulatory elements (REs). OT, optimal transport.

**Supplementary Table 3 | Stage-specific TFs identified by DRAFT and by the SCENIC+-based method of the original study.**

| Developmental stage | Consistent/overlapping findings | SCENIC+-specific findings | DRAFT-specific findings | Main program inferred by DRAFT |
|---|---|---|---|---|
| GSC/Gb (stem cells and gonialblasts) | ovo (germline); esg (stemness); da (bHLH) | klu (self-renewal); Chrac-14 (chromatin); nos (translation); Dp (cell cycle) | TFAM (mitochondria); Mondo/bigmax (metabolic); EcR (ecdysone); Hnf4 (metabolic); cic (growth) | Metabolic/mitochondrial homeostasis as a core GSC/Gb state, with nutrient/endocrine integration (Mondo–EcR/Hnf4) and early state-setting before overt morphological differentiation |
| Spermatogonia (Sp-4 to Sp-16) | foxo (nutrient stress) | fkh (Forkhead); crol (ecdysone); GATAd (GATA); Hr39 (nuclear receptor); Hsf (heat shock); MBD-like (methyl-binding); lov (BTB-ZF) | Xrp1 (stress QC); Mondo (metabolic); Hr4 (ecdysone) ; Mnt (Myc antagonist); cnc (oxidative); chinmo (temporal) | Fitness filtering/QC and metabolic remodeling during the amplification window (Xrp1 with Mondo/Hr4/foxo), consistent with a checkpoint-like program before spermatocyte commitment |
| 1° Sc (primary spermatocytes) | EcR (ecdysone); Hr96 (xenobiotic); usp (RXR); dsx (sex switch); her (bHLH) | Clamp (dosage compensation); Dref (replication); E(bx) (Hox regulation); crc (ATF4); ham (neural) | Rel (NF-κB) | Hormone-gated regulatory remodeling dominated by nuclear-receptor control (EcR/Hr96/usp), consistent with a maturation/priming phase before meiotic execution |
| 2° Sc (secondary spermatocytes) | ken (repressor); dsx (sex switch); Hr96 (xenobiotic); Eip78C (ecdysone); Eip75B (ecdysone) | pros (neural); pdm3 (neural); nub (TF); optix (homeobox); Lim3 (LIM) | cyc (cyclin); disco-r (development); Bap170/D (chromatin) | Transition toward meiotic execution by shutting down growth programs (ken and associated repressors) and activating late-stage reprogramming via hormone/metabolic regulators (Hr96/Eip factors) |

Comparison of stage-specific transcription factors (TFs) identified by DRAFT and by the original study's SCENIC+-based method across developmental stages. Brief functional annotations are given in parentheses. GSC, germline stem cell; Gb, gonialblast; Sp, spermatogonia; Sc, spermatocyte; QC, quality control.

**Supplementary Table 5 | Early germline ranks of Mondo candidate targets across models.**

| Method | *kuz* rank | *mamo* rank |
|---|---|---|
| DRAFT | 13 | 7 |
| DRAFT w/o GRU | 2,617 | 78 |
| Linear + GRU | 146 | 5 |
| Dictys | NA | NA |
| scDoRI | NA | NA |
| SCENIC+ | 51 | 1,967 |

Ranks of *kuz* and *mamo* were evaluated at each of GSC, Gb, Sp-2 and Sp-4. For each method, stage-specific ranks based on Mondo–target scores were combined by their geometric mean and re-ranked to obtain aggregate ranks; lower values indicate higher priority. NA, predictions were unavailable at one or more of the four stages, or the edge weight was zero at all four stages.

**Supplementary Table 6 | Network–expression peak offsets of selected TFs across five DRAFT models.**

| TF | Expression peak | Seed 2024 | Seed 2025 | Seed 2026 | Seed 2027 | Seed 2028 | Earlier / later seeds |
|---|---|---|---|---|---|---|---|
| *Dref* | 2° Sc_1 | Gb (−8) | GSC (−9) | GSC (−9) | Gb (−8) | GSC (−9) | 5 / 0 |
| *Mitf* | GSC | 1° Sc_1 (+6) | 2° Sc_3 (+11) | 2° Sc_3 (+11) | 2° Sc_2 (+10) | 2° Sc_2 (+10) | 0 / 5 |
| *Mondo* | 1° Sc_2 | Sp-2 (−5) | Sp-4 (−4) | Gb (−6) | Sp-2 (−5) | Sp-2 (−5) | 5 / 0 |
| *Max* | 2° Sc_1 | Sp-4 (−6) | Sp-4 (−6) | GSC (−9) | Sp-8 (−5) | Sp-2 (−7) | 5 / 0 |
| *Usf* | 2° Sc_3 | Sp-2 (−9) | Sp-4 (−8) | Gb (−10) | 1° Sc_3 (−3) | Gb (−10) | 5 / 0 |
| *NK7.1* | 2° Sc_1 | Gb (−8) | Gb (−8) | 1° Sc_1 (−3) | Sp-16 (−4) | 2° Sc_4 (+3) | 4 / 1 |
| *Mnt* | 2° Sc_1 | Sp-4 (−6) | Sp-8 (−5) | 2° Sc_6 (+5) | Sp-8 (−5) | 2° Sc_4 (+3) | 3 / 2 |
| *cic* | 1° Sc_2 | Gb (−6) | Sp-4 (−4) | GSC (−7) | Sp-8 (−3) | Sp-2 (−5) | 5 / 0 |
| *bigmax* | 1° Sc_1 | Sp-2 (−4) | Sp-2 (−4) | Sp-16 (−1) | 2° Sc_2 (+4) | 1° Sc_2 (+1) | 3 / 2 |

Parentheses give the network-peak stage minus the expression-peak stage along the ordered 15-stage germline trajectory; negative values indicate an earlier network peak. Network peaks are the maxima of summed outgoing TF–TG scores after Gaussian smoothing ($\sigma$ = 0.5 stage). Earlier/later counts summarize offset direction across the five models and do not apply the conservative ≥3-stage separation criterion based on near-maximum regions. All models use the same observations and regulatory prior; seed 2025 is the manuscript model.

**Supplementary Table 10 | Hyperparameters of DRAFT (unpaired version).**

| Hyperparameter | Value | Description |
|---|---|---|
| in_channels | max_cells | Input dimension for TF/TG node features |
| hidden | 64 | Hidden size for GNN and GRUCell representations |
| out | 64 | Output embedding dimension for TFs and TGs |
| dropout | 0.4 | Dropout rate after GNN layers |
| GNN layers | 2 | Number of heterogeneous GAT layers |
| Optimizer | Adam | Optimizer used for training |
| LR | $2 \times 10^{-4}$ | Learning rate |
| WD | $1 \times 10^{-3}$ | Weight decay (L2 regularization) |
| EPOCHS | 600 | Number of training epochs |
| NEG_RATIO | 1 | Negative sampling ratio |
| Loss | BCE | Applied on sampled positive and negative edges |

hyperparameter settings for implementations of DRAFT (unpaired version)

**Supplementary Table 11 | Hyperparameters of DRAFT (paired version).**

| Hyperparameter | Value | Description |
|---|---|---|
| in_channels | From preprocessing | Input dimension of TF/RE/TG node features |
| hidden_channels | 64 | Hidden size for GNN and GRUCell states |
| out_channels | 64 | Output embedding dimension for TFs, REs and TGs |
| gat_heads | 2 | Number of attention heads in each GATConv |
| dropout_rate | 0.4 | Dropout rate after each GNN block |
| GNN layers | 2 | Two stacked heterogeneous GATConv layers (via HeteroConv) |
| neg_sampling_ratio | 1 | Number of negatives ≈ ratio × number of positives for each edge type |
| num_epochs | 600 | Number of training epochs |
| lr | $5 \times 10^{-4}$ | Learning rate for Adam optimizer |
| weight_decay | $1 \times 10^{-5}$ | L2 regularization strength |
| Optimizer | Adam | Optimizer used for training |
| Loss | BCE | Applied on positive and sampled negative TF–RE and RE–TG pairs |
| lambda_recon | 1 | Trade-off between TF–RE and RE–TG loss |

hyperparameter settings for implementations of DRAFT (paired version)
